# Rational design yields molecular insights on leaf binding of the anchor peptide Macaque Histatin

**DOI:** 10.1101/2022.01.11.475855

**Authors:** Jonas Dittrich, Christin Brethauer, Liudmyla Goncharenko, Jens Bührmann, Viktoria Zeisler-Diehl, Shyam Pariyar, Felix Jakob, Tetiana Kurkina, Lukas Schreiber, Ulrich Schwaneberg, Holger Gohlke

## Abstract

In times of a constantly growing world population and increasing demand for food, sustainable agriculture is crucial. To reduce the amount of applied nutrients, herbicides, and fungicides, the rainfastness of plant protection agents is of pivotal importance. As a result of protective agent wash-off, plant protection is lost, and soils and groundwater are severely polluted. To date, rainfastness of plant protection products is achieved by adding polymeric adjuvants to the agrochemicals. However, polymeric adjuvants will be regarded as microplastics in the future, and environmentally friendly alternatives are needed. Anchor peptides (APs) are promising biobased and biodegradable adhesion promoters. While the adhesion of anchor peptides to artificial surfaces, such as polymers, has already been investigated in theory and experimentally, exploiting the adhesion to biological surfaces remains challenging. The complex nature and composition of biological surfaces such as plant leaf and fruit surfaces complicate the generation of accurate models. Here, we present the first detailed three-layered atomistic model of the surface of apple leaves and use it to compute free energy profiles of the adhesion and desorption of APs to and from that surface. Our model is validated by a novel fluorescence-based MTP assay that mimicks these complex processes and allows quantifying them. For the AP Macaque Histatin, we demonstrate that aromatic and positively charged amino acids are essential for binding to the waxy apple leaf surface. The established protocols should generally be applicable for tailoring the binding properties of APs to biological interfaces.

## 2 Introduction

With the increasing demand for food worldwide, sustainable and innovative plant nutrition and protection methods have become progressively crucial for agricultural production.^1^ An ecologically friendly and tailored application of plant nutrients and protectants may help to reduce the amount of applied substances significantly by improving resistance against rainfall and sunlight and, thus, minimize the ecological footprint. In particular, the rainfastness of foliar applications has been proven to play a major role in its long-term effectiveness and often becomes the limiting factor for the timespan after which reapplication is necessary. Thus, investigating and improving the rainfastness of agrochemicals has been of high interest for a long time.^2-6^ Increased rainfastness is guaranteed by adding adjuvants to the agrochemicals. These adjuvants greatly vary in their chemical nature. Commonly used polymeric adjuvants, however, will be regarded as microplastics in the future.^2, 7-8^ According to the European Chemical Agency, the use of intentionally added microplastics to plant health products will be prohibited within the next five years. Therefore, it is of pivotal importance to identify biodegradable alternatives to ensure future food production.

Anchor peptides (APs) are short amphiphilic peptides with sizes ranging from 20 to 100 amino acids that bind from aqueous solutions to natural^9-10^ and synthetic surfaces^11-14^ including metals^15-16^. Tailor-made APs are thus applicable in many fields, including biotechnology,^17^ catalysis,^18^ nanoparticles,^19^ medicine,^20^ and agriculture^9^.

APs were reported to bind microgel-containers to the plant surface. Such containers can be applied for the long-term controlled release of nutrients, herbicides, and fungicides. APs increase the rainfastness and, hence, reduce the overall amount of chemicals needed for effective plant protection and fertilization.^9^ In another study^10^, APs were fused to an anti-fungal peptide and successfully protected soybean plants against its most severe disease, Asian soybean rust (*Phakopsora pachyrhizi*). Therefore, APs are promising candidates for microplastics-free adhesion-promoting adjuvants to increase rainfastness of agrochemicals.

The antimicrobial AP LCI^21^ shows significant binding to poly(propylene)^13^; the subsequent rational improvement of the binding characteristics to this synthetic material was reported.^11-12^ However, the rational improvement of AP adhesion to biological surfaces remained challenging due to the complex nature and composition of biological surfaces and the diversity of the forces involved.

Here, we developed and validated a workflow for the rational improvement of AP adhesion to specific plant leaves based on a multidisciplinary approach, comprising molecular dynamics (MD) simulations and a tailored fluorescence-based screening experiment. To obtain insights at the atomistic level into the adhesion properties of APs on apple leaves, we generated the, to our knowledge, first three-layered atomistic model of an apple leaf surface consisting of Iβ cellulose, a cutin matrix, and a leaf wax layer and probed AP adhesion to it by molecular simulations. The predictive power of the model was validated by a novel fluorescence-based quantitative assay designed to probe the adhesion of APs towards the leaf surfaces of plants. The generated atomistic models allow investigating not only the binding of APs, but can also aid in scrutinizing the adsorption, penetration, and accumulation^22^ of nutrients, herbicides, pesticides, or fungicides on/through the outer surface of plant leaves. The fluorescence-based assay allows rapidly identifying new potential APs and selecting variants with improved binding properties.

## 3 Materials and Methods

### Chemicals and Materials

All chemicals used in this study were purchased from Carl Roth GmbH (Karlsruhe, Germany), Sigma-Aldrich Corp. (St. Louis, MO and Deisenhofen, Germany), Fluka, (Ulm, Germany), Macherey-Nagel (Düren, Germany) or AppliChem GmbH (Darmstadt, Germany) and had at least analytical-reagent grade purity unless specified. Synthetic genes were obtained from GenScript (Nanjing, China), and oligonucleotides were acquired from Eurofins Scientific SE (Ebersberg, Germany) in salt-free form. Enzymes were obtained from New England Biolabs GmbH (Frankfurt am Main, Germany). Plasmid extraction and polymerase chain reaction (PCR) purification kits were ordered from Macherey-Nagel GmbH & Co. KG (Düren, Germany) and Qiagen GmbH (Hilden, Germany). Black polypropylene microtiter plates (MTPs) were obtained from Greiner Bio-One GmbH (Frickenhausen, Germany). The plasmid pET28a(+) (Novagen, Darmstadt, Germany) was used as the expression vector. The *Escherichia coli* strains DH5α and BL21-Gold (DE3) were purchased from Agilent Technologies (Santa Clara, CA). *E. coli* DH5α was used as cloning host and *E. coli* BL21-Gold (DE3) was used for protein expression.

### Data Acquisition, Model Generation, Assay Development and Model Validation

In the following, we first describe the plant growth conditions and the cuticular wax analysis. Consecutively, this data is used for the *in silico* model generation and molecular simulations for the prediction of binding residues within APs and the binding strength of APs. Finally, the predictions are investigated and validated using a novel quantitative fluorescence-based screening.

### Plant Growth Condition and Leaf Sampling

Stratified apple (*Malus domestica*, cultivar ‘Bittenfelder’) seeds were sown on wet sands for germination. After germination (2-3 leaves stage), seedlings were transplanted in soil pots (1 plant per pot). Seedlings were grown under semi-controlled conditions in the greenhouse to the 12-leaves stage. Then, they were transferred to the field. These apple plants are grown under ambient field environment at the Research Station of Campus Endenich, University of Bonn (50° 73.09’ north latitude, 7° 7.34’ east longitude), and north-west of the city Bonn. Only one plant per pot was allowed to grow after germination to avoid nutrition, soil moisture, and light competition. The plants were grown in well-watered (soil moisture content above 60 %) and nourished soil media (pot with basal-diameter 11 cm, soil mixture: TKS [Brill Typ 5 + sand + perlite, mixing ratio 1:1.2:0.3], water holding capacity: 1.29 kg water kg^-1^ dry mass). Climate data such as global radiation, UVA, rainfall, and photoactive radiation in ambient OD condition were continuously recorded (every minute) at the meteorological station (MWS 9-5 Microprocessor Weather Station, Rheinhardt System-und Messelectronic GmbH, Diessen-Oebermuelhausen, Germany) in the field, whereas plant canopy level temperature plus relative humidity was continuously recorded every 10 minutes by Tinytag data loggers (Tinytag, Gemini Data Loggers, Chichester, UK). During the growth period (18.05-10.07.2017), the global radiation, photoactive radiation, and ultraviolet radiation A in the field were 328 ± 14 W m^-2^ (mean ± standard error (SEM)), 554 ± 20 µmol m^-2^ s^-1^ and 5.0 ± 0.2 W m^-2^, respectively. There were 29 rainfall days with daily cumulative rainfall ranging from 0.11 mm (23.06.2017) to 15.28 mm (10.07.2017), yielding a mean value of 3.5 ± 0.8 mm (SEM) and a median of 1.5 mm rainfall. The canopy level temperature was 23.7 ± 0.5 °C (mean ± SEM), and relative humidity was 61.4 ± 1.4 (mean ± SEM). Fifteen-day-old apple leaves (15 days exposed to field environment) samples were taken for wax analysis from 17^th^-position leaves from the base on 13.06.2017, each from 5 biological replications. The freshly harvested leaves were placed in humid polybags and brought to the lab for wax extraction.

### Quantification of Apple Leaf Wax Components via GC/FID and GC/MS

Surface waxes were extracted from the adaxial leaf side of apple leaves. Waxes were extracted by gently pressing a glass vial containing 5 ml chloroform on the adaxial leaf side for 10 s. Immediately after extraction, samples were spiked with an internal standard (50 µl tetracosane of a chloroform solution of 10 mg in 50 ml; Fluka, Ulm, Germany), enabling the quantification of the individual wax compounds. The chloroform volume was reduced under a gentle stream of nitrogen at 60 °C to an end-volume of 200 µl. Hydroxylic and carboxyl groups of alcohols and acids were transformed into the corresponding trimethylsilyl ethers and -esters by derivatization. Derivatization was done using 20 µl *N,O*-bis(trimethylsilyl)-trifluoroacetamid (BSTFA; Macherey-Nagel, Düren, Germany) and 20 µl pyridine (Sigma Aldrich Corp., Deisenhofen, Germany) for 45 min at 70 °C. 1 µl of each sample was analyzed by on-column injection and a gas chromatography equipped with flame ionization detection (GC-FID; CG-Hewlett Packard 5890 series H, Hewlett-Packard, Palo Alto, CA, USA, column-type: 30 m DB-1 i.d. 0.32 mm, film 0.2 µm; J&W Scientific, Folsom, CA, USA). For identification of the individual wax components (i.e., fatty acids, alcohols), again 1 µl of the samples were analyzed by GC-MS (Quadrupole mass selective detector HP 5971, Hewlett-Packard, Palo Alto, CA, USA). All wax molecules, including the fatty acids, were quantified based on the amount of internal standard via GC-FID analysis. Identification of the single compounds was made using a homemade wax database and by comparing the obtained fragmentation pattern with known substances. Calculations were performed for every single component individually. The raw data is provided in Table S1.

### Computational Methods

To rationally improve the adhesion of the tested APs to the surface of an apple leaf, we generated an atomistic model of an apple leaf surface and performed (steered) molecular dynamics simulations of the ad- and desorption of the tested APs to/from this model.

### Generation of an All-Atom Model of a Leaf Surface

The outer part of the leaf surface consists of a cuticular polyester matrix of inter-esterified ω-hydroxy acids, impregnated with cuticular waxes, covered with cuticular waxes,^23^ and located atop of a polysaccharide cell wall.^24^ We used the Cellulose-Builder^25^ to generate a crystalline sheet of Iβ cellulose, consisting of three layers of 15 chains each with 15 β(1→4)-linked D-glucopyranose moieties per chain, yielding a sheet of 155 Å x 121 Å x 8 Å size (Figure 1A) that serves as a rigid surface for our cutin model. Force field parameters for the β(1→4)-linked D-glucopyranose units and the terminal hydroxyl groups were taken from GLYCAM06.^26^

**Figure 1.**
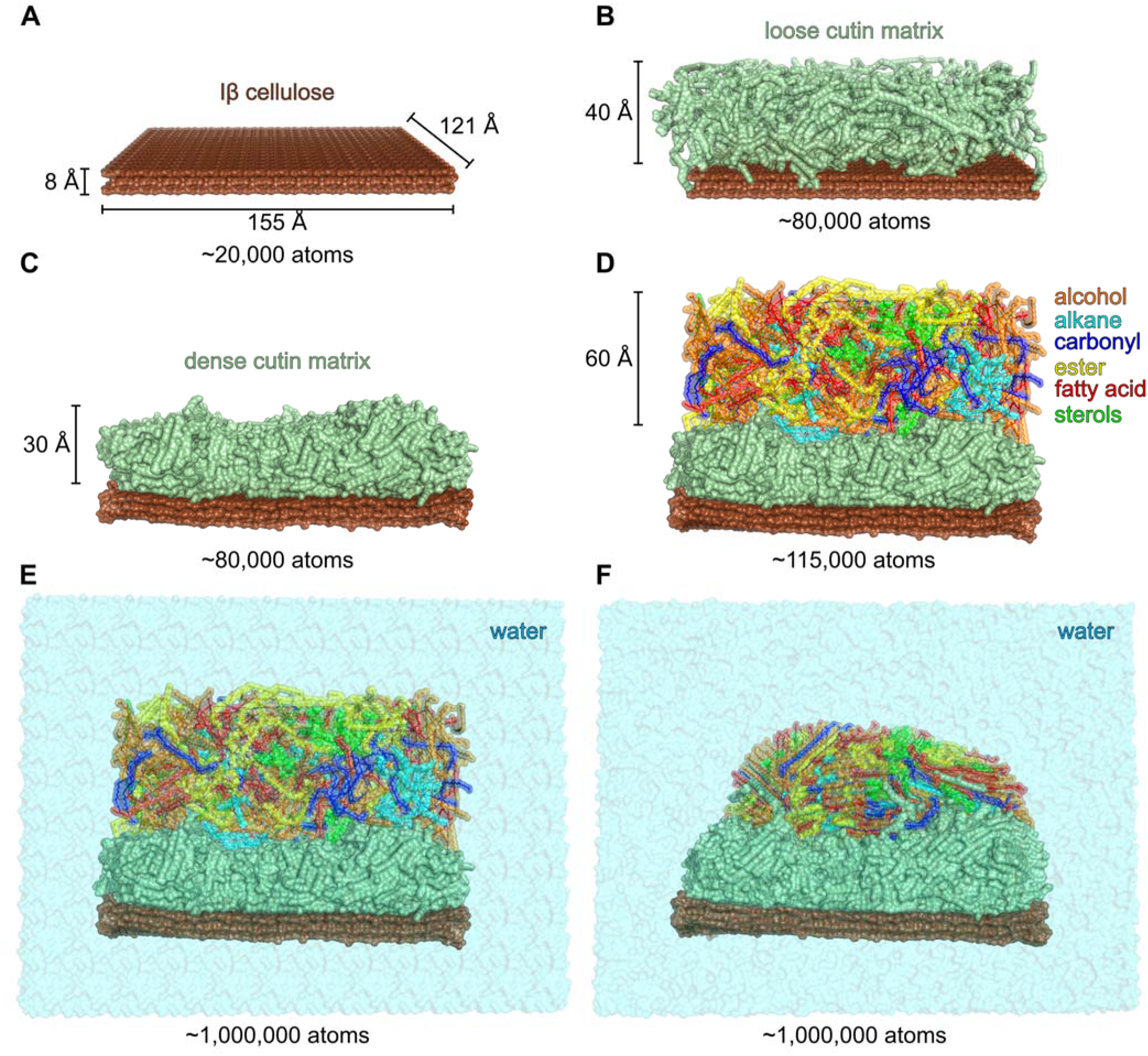
Stepwise creation of an atomistic leaf surface model. **A)** Three-layered crystalline Iβ cellulose. **B)** Loosely packed polyester matrix of cross-linked 10,18-dihydroxyoctadecanoic acid located above the cellulose sheet. **C)** Compacted cutin layer atop the cellulose sheet after 25 ns of NVT MD simulations. **D)** Randomly placed wax components above the compact cutin layer. **E)** Solvated system. **F)** Snapshot obtained after 100 ns of NVT MD simulations of the complete model.

Cutin is a waxy polymer composed of ω-hydroxy acids (mostly long-chain (16- or 18-carbon) fatty acids) and their derivatives, which are interlinked via ester bonds, forming a polyester polymer of indeterminate size. Its composition varies depending on the plant.^27^ As a representative fatty acid, we chose 10,18-dihydroxyoctadecanoic acid. To generate building blocks for all possible ester cross-links, all possible 2^3^ structures differentially methylated at all hydroxyl groups were generated in Schrödinger’s MAESTRO software suit.^28^ In subsequent preparation steps, methyl caps were removed, connection records were added instead, and the structures were saved in an Amber library file (.lib), allowing one to generate custom polyester matrixes. We generated several linear polyesters, ranging from dimers to hexamers, and branched structures with up to 17 residues. The polyesters were packed into a rectangular box (160 Å x 120 Å x 80 Å) atop the cellulose using PACKMOL,^29^ thereby keeping a minimal distance of at least 2 Å between all structures. This loosely packed cutin matrix (Figure 1B) was solvated using TIP3P water^30^ and energy-minimized and thermalized. Then, MD simulations of 25 ns length under NVT conditions (see next section for details) were performed to compact the matrix (Figure 1C). The water from the resulting cellulose/cutin model is stripped, and the system is used as a starting structure for all consecutive MD simulations performed to generate the complete atomistic model of the leaf wax on a cutin surface. Note that the thickness of the cutin layer may vary considerably in nature.^23^ However, as we are focusing on interactions of the peptides with the leaf surface, our polyester layer merely serves to mimic the outer part of the cutin layer in order to adequately model interactions between the leaf surface and the wax components.

The composition of the wax is based on experimental data obtained by GC/FID and GC/MS of an apple leaf wax/chloroform extract (see “Quantification of apple leaf wax components via GC/FID and GC/MS”). To keep our computations tractable, we used 1/45 of the experimentally determined amount of wax components per surface area of the cellulose/cutin system. PACKMOL^29^ was used to pack the wax components into a rectangular box atop the cellulose/cutin system (Figure 1D). Ten initial starting structures were created by selecting different random seeds for the packing process of the wax components. The resulting systems were again solvated (Figure 1E) with TIP3P^30^ water such that the distance between the boundary of the box and the closest solute atom was at least 20 Å. MD simulations of 100 ns length in the NVT ensemble were then performed (Figure 1F), as described in the following section after minimization and thermalization. Atomic charges of cutin and wax components were determined using the AM1-BCC method^31-32^ as implemented in antechamber.^33^ Force field parameters were taken from the GAFF2 force field.^34^ Na^+^ counterions were used to balance the charges of the deprotonated fatty acids. The solvated system comprises more than 1,000,000 atoms. At the time the model was constructed, AMBER did not have the option to handle covalent bonds across periodic boundaries. Therefore, we were limited to a finite sheet of cellulose which has to be restrained during simulations (see Molecular Dynamics Simulations) and the padding of the system with solvent molecules.

The coordinates of the ten generated leaf surface models are provided in PDB format and Amber .inpcrd. and .prmtop files in the Supporting Information.

### AP/Leaf Systems

Peptide structures were taken from the Protein Data Bank (PDB)^35^ when available (name, PDB ID: LCI, 2B9K; Magainin, 2MAG; Plantaricin A, 1YTR; Pleurocidin, 1Z64). If a PDB entry contained an ensemble of structures, the first structure was taken, and if the protein was present as a multimer, the first subunit was used. We employed TopModel^36^, a meta-method for protein structure prediction using top-down consensus and deep neural networks, to generate a structural model for Macaque Histatin (MacHis). Ten systems for the simulation of the adhesion for each peptide were created by placing three APs randomly above the ten different leaf wax replicas using PACKMOL, while keeping a minimum distance of at least 5 Å between the APs as well as to the wax surface. Na^+^/Cl^-^ counterions were added to neutralize the charges. Thus, for all peptides, ten independent MD simulations of 250 ns length each were performed, using differently packed leaf wax models for each system. To probe the influence of the initial conformation of the leaf wax surface on the simulation, we simulated ten replicas of LCI for each of the ten initial leaf wax models for 250 ns, thus, in total 100 runs.

### Molecular Dynamics Simulations

MD simulations were carried out with the Amber18 suite of programs^37-38^ using the GPU-accelerated CUDA version of PMEMD^39-40^. We applied the ff14SB^41^ (for the peptides), GLYCAM06^26^ (for Iβ cellulose), and GAFF2 (for cutin and wax components) force fields^34^ in all simulations. The structures were solvated in a box of TIP3P^30^ water such that the distance between the boundary of the box and the closest solute atom was at least 20 Å. Periodic boundary conditions were applied using the particle mesh Ewald (PME) method^42^ to treat long-range electrostatic interactions. Bond lengths involving bonds to hydrogen atoms were constrained by the SHAKE^43^ algorithm. The time step for all MD simulations was 2 fs, and a direct-space non-bonded cutoff of 8 Å was applied. First, the solvent was minimized for 250 steps by using the steepest descent method followed by conjugate gradient minimization of 50 steps. Subsequently, the same approach was used to minimize the entire system. Afterwards, the system was heated from 0 K to 100 K using canonical ensemble (NVT) MD simulations, and from 100 K to 293 K using isobaric MD simulations. The solvent density was adjusted to 0.97 g cm^-3^ using isothermal-isobaric ensemble (NPT) MD simulations. Positional restraints applied during thermalization were reduced in a stepwise manner over 50 ps, followed by 50 ps of unrestrained canonical ensemble (NVT) MD simulations at 293 K with a time constant of 2 ps for heat bath coupling with the Berendsen thermostat^44^. Production MD simulations for the leaf model only and for the system comprising the APs were run for 100 ns and 250 ns length, respectively, using 10 ps for heat bath coupling^44^. Coordinates were saved at 100 ps intervals. To prevent twisting of the cellulose layer, positional restraints with a force constant of 1 kcal mol^-1^ Å^-2^ were applied to the glucopyranose moieties throughout the production simulations.

### Estimation of the PMF for AP Desorption by Adaptive Steered Molecular Dynamics Simulations

To estimate the work needed to release the AP from the wax surface, we performed adaptive-steered molecular dynamics (ASMD) simulations.^45-46^ Using Jarzynski’s equality,^47^ the nonequilibrium work *W*_A→B_ performed on the system in a steered MD simulation can be related to the free energy difference Δ*F* = *F*_B_ – *F*_A_ between state A and B^48^ as depicted in eq. 1 and eq. 2 (*T*: absolute temperature; *k*_B_: Boltzmann constant):

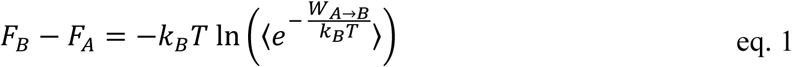

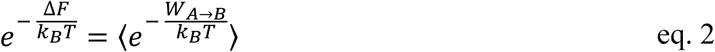

In steered MD simulations (SMDs), a steering force is applied along the desired reaction coordinate *ξ*(*r*) at a constant velocity *ν* to explore the system.^45, 48^ The system’s original Hamiltonian *H*(*r, p*) is extended by a guiding potential *h*_A_(*r*), with the spring constant *k* and the time-dependent perturbation *λ* = *λ*(*t*) (eq. 3), yielding the total Hamiltonian 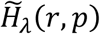 (eq. 4)^48^:

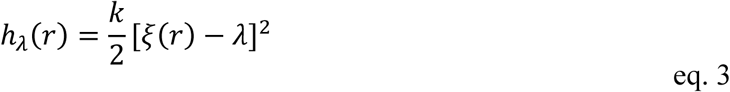

with

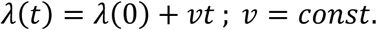

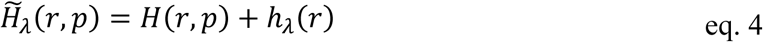

When applying Jarzyski’s equality to the 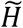 system, we obtain eq. 5:

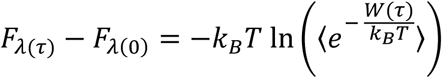

with

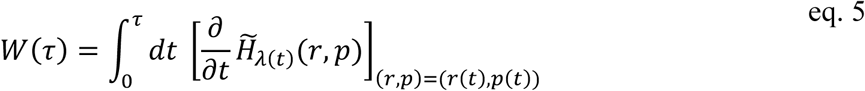

where *F*_A_ is the Helmholtz free energy of the 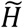 system and *W*(*τ*) is the work done during the time interval [0, *τ*] calculated for each trajectory (*r*(*t*), *p*(*t*)).

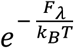 can be expressed in terms of the Helmholtz free energy profile (potential of mean force) *PMF*(*ξ*) along *ξ*, with the highest contribution of the integral coming from the region around *ξ* = *λ* for large *k*, known as stiff-spring approximation.^48^ Taking the Taylor series of 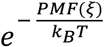 about *λ*, then, allows calculating the PMF from the leading order, yielding *PMF*(*λ*) = *F*_A_.^48^ The latter can be calculated from eq. 5, resulting in *PMF* as a function of the distance *d* from the leaf surface plus a constant *const* = *F*_*λ*(0)_ according to eq. 6:

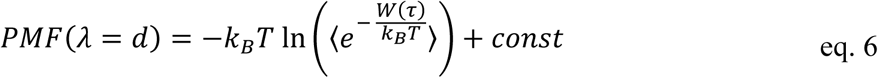

In general, a high number of simulations must be performed to converge the average over all realizations of an external process that takes the system from the equilibrium state A to a new, generally nonequilibrium state B and, thus, obtain a converged PMF.^48^ ASMD tackles this problem by dividing the reaction path into smaller segments called stages.^49^ Within these stages, the trajectory closest to the Jarzynski average is determined, and the final state of this trajectory is used as starting point for the consecutive stage.^50^ Thus, trajectories contributing little to the overall PMF are disregarded in the next stage, that way reducing the total number of simulations to be performed.

Here, we performed ASMD simulations with a pulling velocity of *ν* = 1 Å ns^-1^ along the surface normal of the leaf model, and only the work applied along this axis is determined. We employed a uniform force constant of *k* = 100 kcal mol^-1^ Å^-2^. For each stage, 25 replicas are simulated for 2 ns each, resulting in an increase of 2 Å in the distance between the center of mass of the AP and the leaf surface per stage. As the amount of required simulation time is considerable for a system comprising more than 1,000,000 atoms (∼6-8 hours on state-of-the-art GPUs for 1 ns of simulation time), we limited the tested APs to MacHis, LCI, Plantaricin A, Magainin, and Pleurocidin. These peptides cover the range from strong adhesion (MacHis, LCI) over medium adhesion (Plantaricin A) to weak/no significant adhesion (Magainin, Pleurocidin) as determined from qualitative experiments on the apple. For each AP, one system out of the ten replica is chosen for ASMD simulations, where one of the three APs present in the system is adsorbed approximately at the center of the wax surface and not interfering with the remaining APs.

### Identification of Residues Contributing to Binding

To identify residues important for the adhesion of the AP to the leaf surface, we calculated all contacts of the peptides to the wax components within a distance cutoff of 7 Å for each replica. Based on the identified interactions, MacHis variants were generated by substituting the residues showing the highest number of contacts with alanine. As a negative control, alanine variants of residues showing less frequent contacts were generated (see experimental approach). The geometric analyses of the trajectories were performed with CPPTRAJ.^51^

### Assay Development and Model Validation

To quantify the AP’s binding strength to the cuticular wax and validate the computational predictions, we developed a fluorescence-based MTP assay to screen the APs and AP variants rapidly. In the following, we describe the developed assay and the generation of the AP fusion constructs.

### Wax Extraction and Coating of MTPs

For cuticular wax extraction, apple leaves (*Malus domestica*, cultivar ‘Pinova’, 200 leaves) were immersed in pure chloroform for 10 sec. The remaining cuticular wax-chloroform mixture was filtered (cellulose folded filter paper; 4-12 µm pore size, M&N 615 1/4, Ø 185 mm, Macherey-Nagel GmbH & Co. KG), and chloroform was evaporated (rotary evaporator; 500 mbar to 250 mbar, 49-50 °C water bath, IKA^®^-Werke GmbH & CO. KG) (Figure 2A). For the MTP wax coating, the wax-chloroform mixture was added to each well (50 µg/cm^2^ for apple wax; 50 µl per well), and chloroform was evaporated completely (RT, 16 h) (Figure 2B). Wax-coated MTPs were subsequently used for the characterization of AP binding strength.

**Figure 2.**
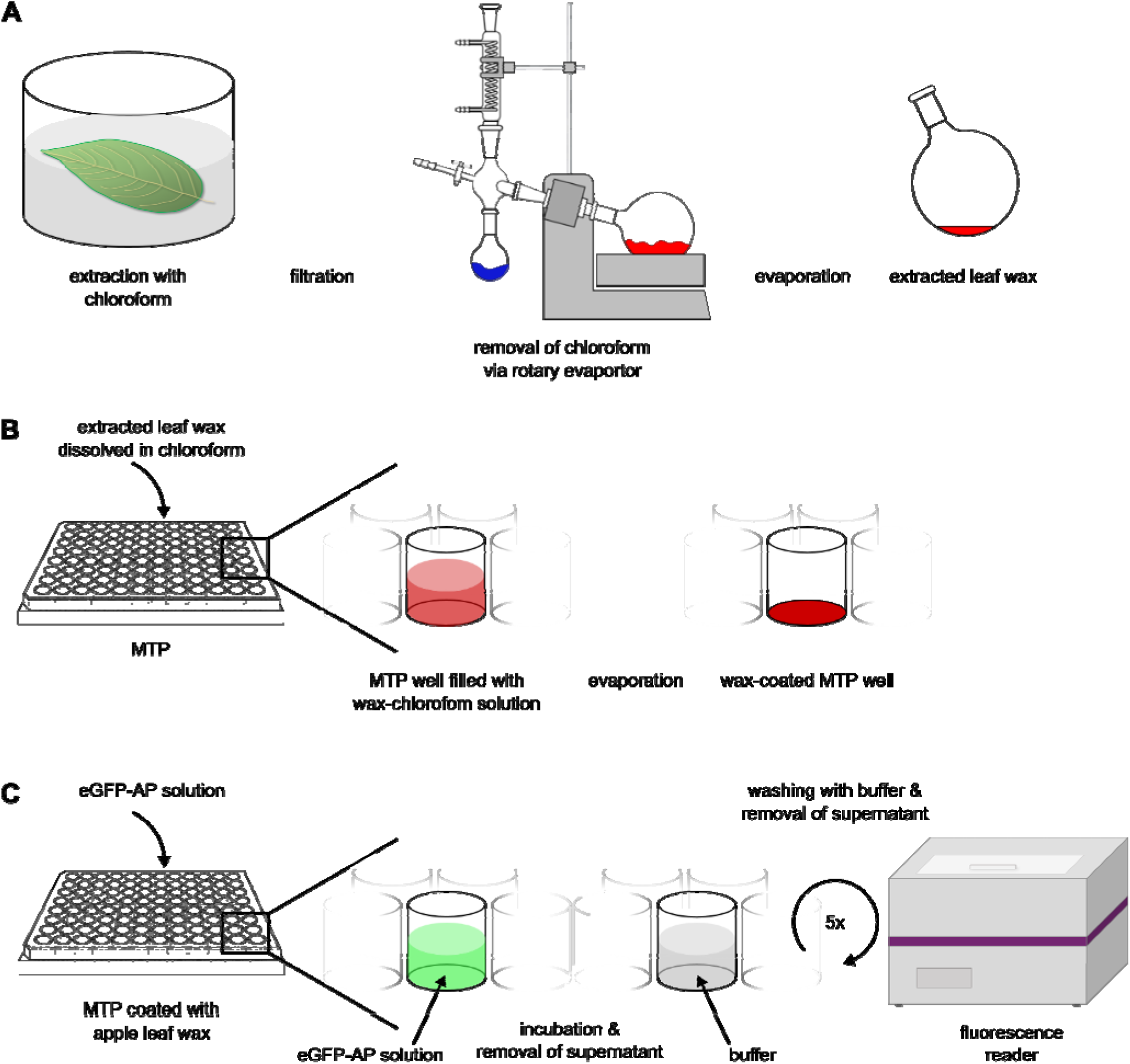
Generation of an MTP-based assay for evaluating the adsorption of APs towards surface wax. **A)** Surface wax extraction of an apple leaf using chloroform. The extracted wax can be stored and (re)dissolved for coating. **B)** Wax coating of an MTP well. **C)** Procedure of the binding assay describing the application of the eGFP-AP solution and consecutive washing steps. The remaining fluorescence is detected using a fluorescence reader.

### Generation of eGFP-AP Fusion Constructs

The synthetic genes of LCI (UniProt ID: P82243), MacHis (UniProt ID: P34084), Magainin (UniProt ID: P11006), Plantaricin A (PlnA; UniProt ID: P80214), and Pleurocidin (UniProt ID: P81941) were codon-optimized for *E. coli* and synthesized by GenScript (Nanjing, China) (Table S2). All synthetic genes contained a stiff spacer helix (17 amino acids; AEAAAKEAAAKEAAAKA)^52^ followed downstream by a TEV cleavage site (7 amino acids: ENLYFQG)^53^ at the 5’-end of the AP gene. APs were cloned in the pET28a(+)::eGFP backbone applying “sequence-independent phosphorothioate-based ligase-independent gene cloning” (PLICing)^54^ as previously described.^13^ For the performed binding study, the TEV cleavage site was removed. Primers were designed (Table S3), and a two-step PCR amplification was performed under the following conditions: pre-PCR for single-primer extension ((98 °C, 2 min; one cycle), (98 °C, 15 s/ 55 °C, 15 s/72°C, 4 min; 6 cycles)) followed by a final elongation step (72 °C, 10 min; one cycle) and a PCR for efficient recombination ((98 °C, 2 min; one cycle), (98 °C, 15 s/ 55 °C, 15 s/72 °C, 4 min; 15 cycles)) followed by a final elongation step (72 °C, 10 min; one cycle). All generated constructs were digested (20 U *Dpn1*; 2 h, 37 °C) and purified using PCR clean-up kit (Macherey-Nagel). As eGFP-control, the construct pET28a(+)::eGFP-17xHelix was generated. All generated constructs were transformed in electrocompetent *E. coli* DH5α and BL21-Gold (DE3) cells. Successful cloning was confirmed by sequencing (Eurofins Genomics GmbH, Ebersberg, Germany).

### Generation of eGFP-MacHis Variants

Amino acid substitutions were introduced into the eGFP-MacHis system by side-directed mutagenesis using the pET28a::eGFP-17xHelix-TEV-MacHis template and the primer pairs shown in Table S1 and Table S2 using a two-step PCR. The PCR conditions were: pre-PCR for single-primer extension ((98 °C, 2 min; one cycle), (98 °C, 15 s/55 °C, 15 s/72 °C, 4 min; 6 cycles)) followed by a final elongation step (72 °C, 10 min; one cycle) and a protocol for efficient introduction of point mutations ((98 °C, 2 min; one cycle), (98 °C, 15 s/ 55°C, 15 s/72 °C, 4 min; 15 cycles)) followed by a final elongation step (72 °C, 10 min; one cycle). All generated constructs were digested (20 U *Dpn1*; 2 h, 37 °C), purified using PCR clean-up kit (Macherey-Nagel), and transformed in electrocompetent *E. coli* DH5α and BL21-Gold (DE3) cells. Successful construction and cloning were confirmed by sequencing (Eurofins Genomics GmbH, Ebersberg, Germany).

### Production of eGFP-AP Fusion Constructs

eGFP-control, eGFP-LCI, eGFP-MacHis, eGFP-Magainin, eGFP-PlnA, eGFP-Pleurocidin, and eGFP-MacHis variants were expressed in *E. coli* BL21-Gold (DE3) cells and produced in Erlenmeyer flasks as published before.^13^ Cultivation was performed for 24 h (20 °C, 900 rpm, 70 % humidity; Multitron Pro; Infors AG). Cells were harvested by centrifugation (3200 g, 40 min, 4 °C; Eppendorf centrifuge 5810 R, Eppendorf AG, Hamburg, Germany), and the cell pellets were stored (−20 °C). The obtained cell pellets were suspended in tris(hydroxymethyl)-aminomethane (Tris/HCl) buffer (10 mL; pH 8.0, 50 mM) and disrupted by sonication on ice (2 × 3 min, interval 30 s, 70 % amplitude). By centrifugation (3200 g, 30 min, 4 °C; Eppendorf centrifuge 5810 R), soluble proteins were separated from cell fragments and insoluble proteins. The supernatant was filtered through a 0.45 mm cellulose-acetate filter (GE Healthcare, Little Chalfont, UK) and subsequently used for further purification.

### Purification of eGFP-AP Fusion Constructs

The eGFP-AP fusion proteins and the eGFP-control were purified using the N-terminal His_6_-Tag and fast protein liquid chromatography (ÄKTAprime, GE Healthcare) with a prepacked ion affinity chromatography column (Ni-Sepharose 6 Fast Flow, 5 mL, GE Healthcare). Samples were eluted with imidazole, desalted by dialysis against Tris/HCl buffer (50 mM, pH 8.0; used membrane: Spectra/Por®4, Spectrum Inc., Breda, The Netherlands) and concentrated using ultrafiltration (Amicon® Ultra 15 mL Centrifugal Filters, Merck KGaA). Protein concentrations were determined with the BCA protein assay kit (Novagen, Merck KGaA), and protein homogeneity was analyzed by sodium dodecyl sulfate-polyacrylamide gel electrophoresis (SDS-PAGE; 5 % stacking gel, 12 % separation gel).

### Characterization of eGFP-AP Binding Strength to Apple Leaf Wax

To analyze the binding strength of the different eGFP-APs and eGFP-MacHis variants towards apple leaf wax, a 96-well MTP-based assay was established (Figure 2). In the immobilization step, eGFP-APs (4 µM and 10 µM for eGFP-APs and eGFP-MacHis variants, respectively; supplemented to PBS buffer pH 7.4, 100 µL per well) were transferred to a black wax-coated MTP (PP, flat bottom, wax coating described above) and incubated (10 min, RT, 600 rpm, MTP shaker, TiMix5, Edmund Bühler GmbH, Hechingen, Germany). In five subsequent washing steps, the MTP wells were washed with PBS buffer (100 μl/well; 5 min, RT, 600 rpm) to remove non-specific as well as weak binding peptides and select strong binding eGFP-APs. After removal of the liquid and desorbed peptides, the fluorescence of the bound eGFP-APs was measured directly on the wax-coated MTP surface with the 96-well MTP reader FLUOstar Omega (BMG LABTECH GmbH, Ortenberg, Germany; excitation (exc.) 485 nm, emission (em.) 520 nm, gain 1,000, 35 reads/well). The obtained fluorescence is normalized by the fluorescence of the respective “dry” well after the initial treatment and removal of the supernatant to account for varying concentrations. Finally, a baseline correction is performed to account for the fluorescence of the pure eGFP.

### Immobilization of eGFP-APs to Apple Leaves

Apple leaves were cleaned with water and cell-free extracts containing eGFP-APs (50 µL) were added to the apple leaves and incubated at room temperature. All non-specific or weak binding peptides were removed in a subsequent washing step (50 mM Tris-HCl buffer pH 8.0, 1 mL, 3 min incubation). Immobilization of the APs on apple leaves was confirmed by detection of the fluorescent fusion partner eGFP and visualized by confocal microscopy (Leica Microsystems GmbH, Ex: 335 nm, Em: 454 nm, 405 diode laser).

## 4 Results and Discussion

Here, we present our multidisciplinary workflow (Figure 3), ranging from apple leaf sample selection in the field over surface wax extraction, analysis, and MTP assay preparation on the laboratory scale to molecular modeling and simulations and back to the laboratory scale for the screening of the identified APs and AP variants. Thus, in the following, we will first present our findings with regard to the adaxial surface wax composition. Second, we will show the results of unbiased MD simulations involving the atomistic apple leaf surface model to identify residues of MacHis essential for binding to that surface and the experimental validation. Third, we show that our model can qualitatively predict the binding strength of structurally different APs, as the results match the ones obtained from the fluorescence assay.W

**Figure 3.**
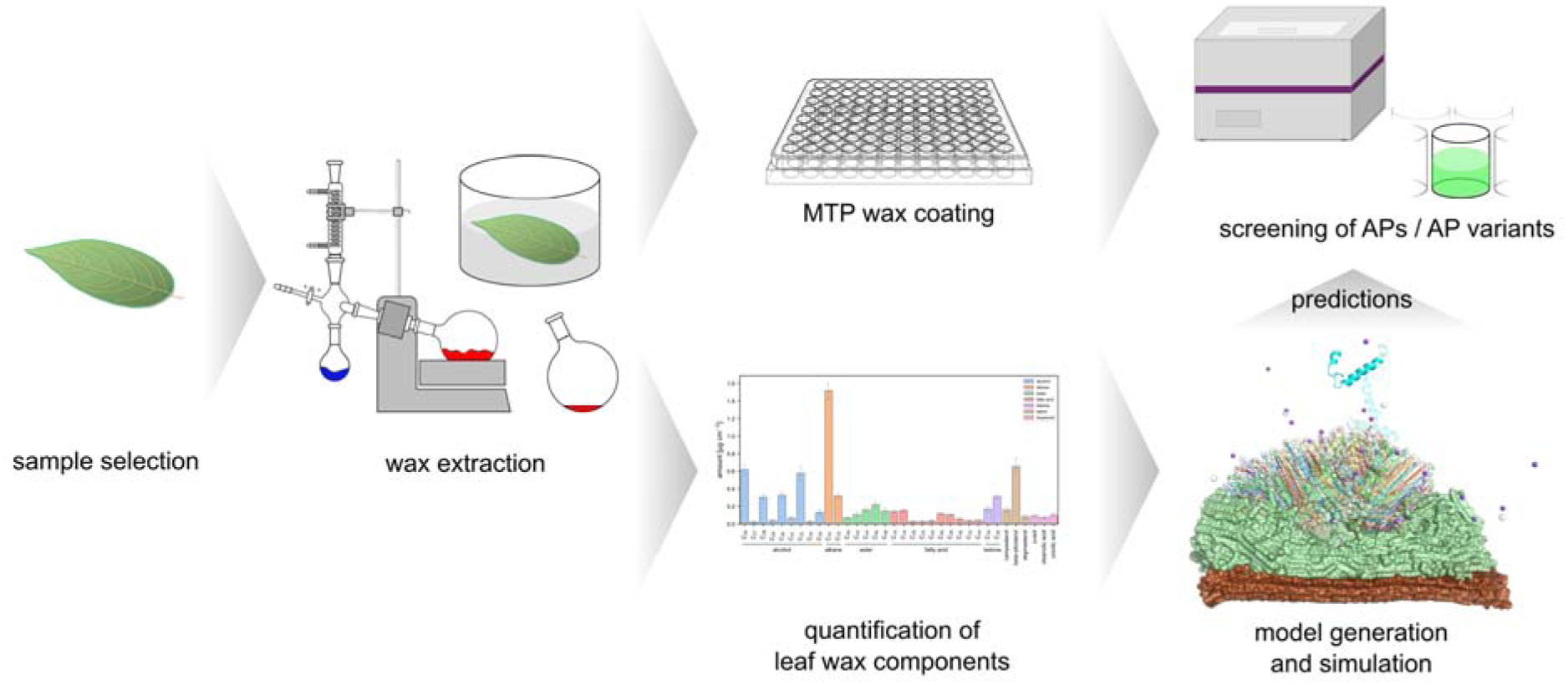
Multidisciplinary workflow for the rational improvement of the binding strength of APs towards apple leaves.

### Apple Leaf Wax Composition

Analytical investigations of the adaxial leaf side of *Malus domestica*, cultivar ‘Bittenfelder’ using GC/FID and GC/MS indicate that the C_31_ alkane is the most prominent compound. Besides alkanes, primary alcohols (chain length C_26_ to C_34_), esters (chain length C_40_ to C_48_), primary acids (chain length C_16_ to C_34_), and ketones were the dominating linear long-chain aliphatic wax compounds. In addition, triterpenoids representing typical wax compounds for species of the plant family *Rosaceae* were found (Figure 4, Tabel S3). Sterols were also identified in the wax extracts, however, since they are typical cell membrane components, it cannot be excluded that they partially also originated from the leaf interior. Our findings are in line with published data^55^ on the adaxial apple leaf wax composition, where alkanes, alcohols, esters, and acids of similar size and triterpenoids were described as prominent wax compounds, whereas steroids were missing in this study. The complex composition of the leaf wax does not allow drawing conclusions about the precise physicochemical properties of the wax layer and potential interactions with the APs, requiring MD simulations for further investigations. Our workflow for the extraction of surface wax from different kinds of leaves, e.g., different species, different growing conditions, and different positions on the plant, allows a standardized determination of the wax composition, and the resulting knowledge about the composition can be used to adapt our atomistic model to reflect the complex variety of surface waxes.

**Figure 4.**
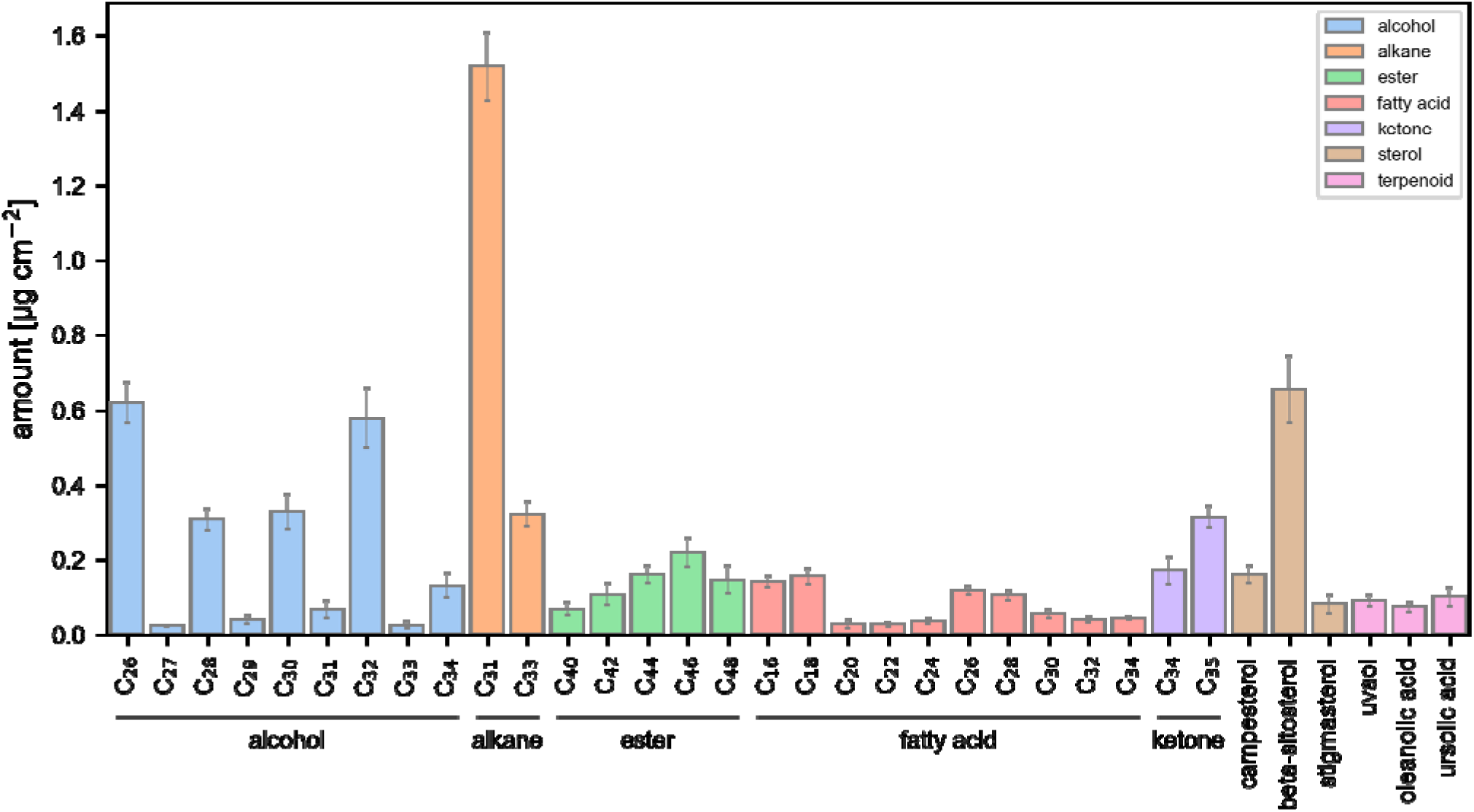
Composition of apple (*Malus domestica*, cultivar ‘Bittenfelder’) adaxial leaf wax obtained via GC/FID and GC/MS. The bars depict the mean (*n* = 5) amount of the respective wax component, and the error bars depict the standard error of the mean.

### Generating an Atomistic Model of an Apple Leaf Surface

Using the determined composition of the apple leaf surface waxes, we tailored the wax layer of our three-layer leaf surface model to represent an apple leaf’s outer surface adequately. As the generation of the model is a multi-step process (Figure 1), a high number of alternative and independent conformations of the system results. Here, we used one template of the cellulose-cutin model (Figure 1C) for the generation of ten systems exhibiting differently packed wax layers to promote the versatility of the cuticular wax layer and account for natural changes within it. In principle, the composition of the wax layer can be adjusted to experimental data to reflect different kinds of plant leaves and growing conditions. Besides allowing to identify key interactions of APs (see below), the model should be applicable to investigate deposition and penetration processes of agrochemicals through the different layers, potentially leading to accumulation in plants.^56-58^ The generated atomistic model provides a detailed description of the outer leaf surface and is a platform for further coarse-graining^59-60^ to speed up computational evaluation in the future, when a simplified topology is sufficient. Here, we note that the generated models show a noticeable accumulation of the leaf wax components. Due to the solvation (at least 20 Å between the boundary of the box and the closest solute atom) during the preparation of the three-layered model (see Generation of an All-Atom Model of a Leaf Surface in the Methods section), the wax components accumulate in the center of the cutin plain to minimize the hydrophobic area exposed to the solvent.

### Identification of Preferred Leaf Wax-Binding Residues within MacHis

We identified the preferred binding residues of MacHis from the ten unbiased MD simulations of 250 ns length, each containing a leaf model and three AP moieties. We evaluated all contacts of the MacHis to the wax components within a distance of 7 Å and considered residues to preferentially bind to the leaf wax if they form > 5 % of all interactions (Figure 5A). The secondary structure of MacHis is predicted^36^ to be predominantly an α-helix with a small loop region between residues 7 and 11. The MD simulations reveal that MacHis preferentially binds to the cuticular wax with one side of the α-helix (Figure 5B). Residues located at positions 6, 10, 16, and 20 show the highest number of contacts (Figure 5A). These positions were selected for alanine scanning in combination with the high-throughput MTP assay to determine the importance of these residues for binding to the cuticular wax. The contact analysis for the remaining four investigated APs is shown in Figure S1.

**Figure 5.**
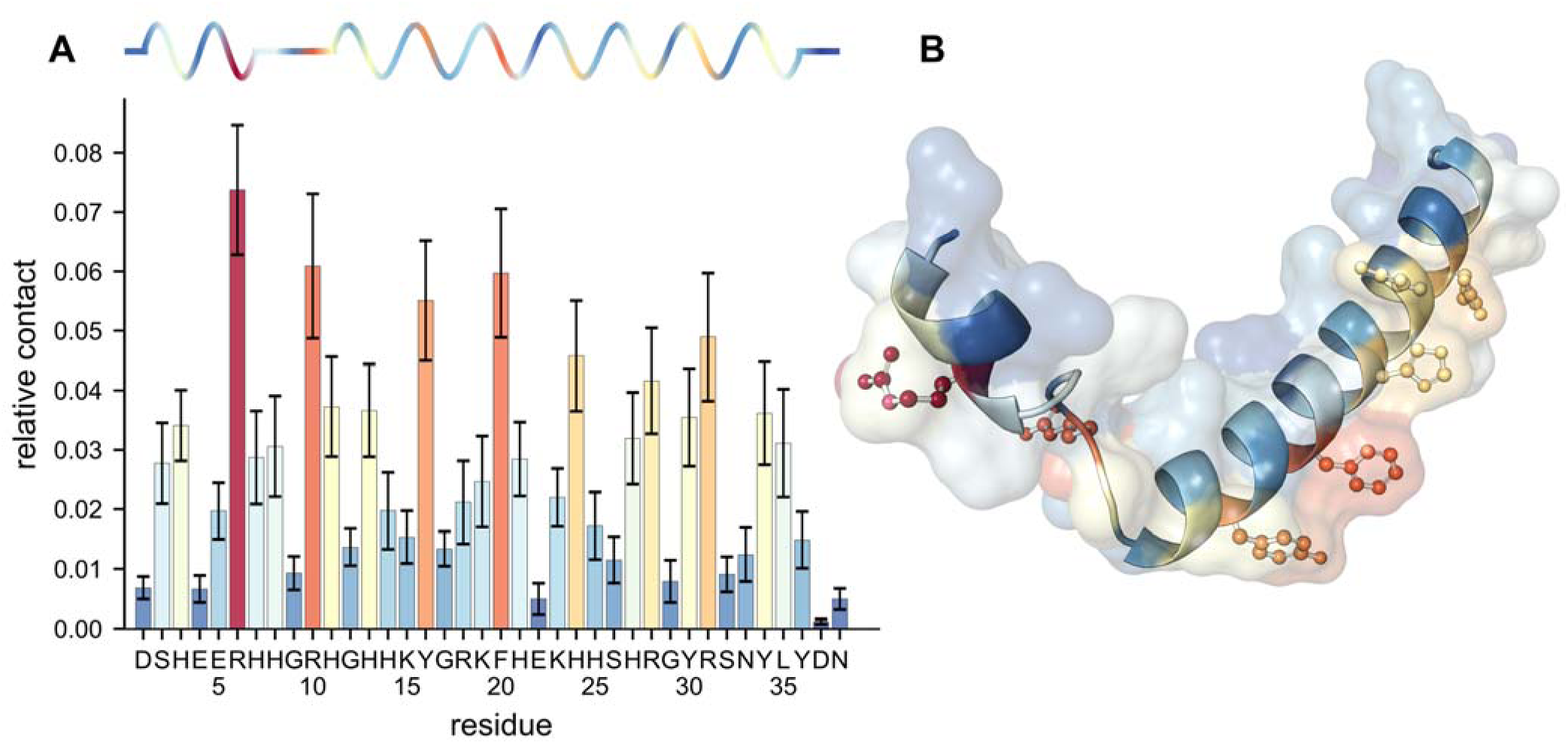
**A)** Residue-wise relative contacts of MacHis with the wax molecules during 10 × 250 ns of MD simulations of AP adsorption. The secondary structure, as determined by DSSP,^61^ is indicated on the top. The color code relates to that shown in panel B. Error bars denote the SEM. **B)** Homology model of MacHis colored according to the relative number of contacts a residue forms within 7 Å with the wax molecules; sidechains of residues with a relative contact > 0.04 are depicted as sticks and spheres.

In order to probe the dependency from the sample size, i.e., the number of observed AP-wax interactions, on the identification of preferred binding residues, we performed bootstrapping analyses on the MacHis (Figure S2) and LCI (Figure S3) simulation data. When using a sample size of 15 APs, i.e., approximately half of the MacHis set, in 71 % (17 %) of all cases, it is possible to identify two (three) out of the three residues identified using the complete data set. When considering the five best binding residues, in more than half of the cases (53 %), it is still possible to identify four out of the five residues (in 13 % of all cases, all five residues are identified), using half of the data set. In comparison, the bootstrapping analysis of the exhaustive simulations using LCI reveals that, with our setup (10 replica à 3 APs), one can identify at least two out of the top three (three out of the top five) residues identified using a ten times larger dataset in ∼80 % (∼90 %) of the cases, suggesting that the chosen setup (10 replica à 3 APs) provides a reasonable balance between accuracy and computational demand.

Finally, a simulation time of 250 ns for adsorption proved to be sufficient, as upon binding to the wax layer, which is observed within less than 100 ns for the majority of all MacHis moieties, the binding pose of MacHis is stable for the remaining simulation time as indicated by only minor changes compared to the previous frame (<RMSD_previous,100-250 ns_> = 2.16 ± 0.04 Å, Figure S4).

### Alanine Scanning of the Identified Residues of MacHis Reveals their Importance for Binding to the Surface Wax

As a general trend, positively charged and aromatic amino acids (R6, R10, Y16, F20) were identified as preferred binding residues interacting with the wax layer. Amino acids at the four binding residues with the highest relative contact were exchanged to alanine (Figure 5); the generated variants are MH_1_ (R6A), MH_2_ (R10A), MH_3_ (Y16A), MH_4_ (F20A), MH_5_ (R6A/R10A), MH_7_ (R6A/R10A/F20A), and MH_8_ (R6A/R10A/Y16A/F20A). In addition, one eGFP-MacHis variant, MH_6_ (K15A), with an amino acid substitution at a residue with few relative contacts to the apple leaf wax was generated as a negative control. Overall, seven eGFP-MacHis variants were generated by site-directed mutagenesis.

The APs’ binding strength to apple leave wax was quantified using the eGFP fluorescence. Through washing steps, peptides and proteins with weak interactions to the cuticular wax are removed, whereas peptides with higher affinity remain on the surface. Therefore, the binding strength of eGFP, eGFP-MacHis wild-type (MH_WT_), and the eGFP-MacHis variants (MH_1-8_) correlates with the determined fluorescence after washing. The remaining fluorescence of eGFP, MH_WT_, and MacHis variants after five washing steps is depicted in Figure 6 (raw data is provided in Tables S4-S6, the remaining fluorescence after two washing steps is depicted in Figure S5).

**Figure 6.**
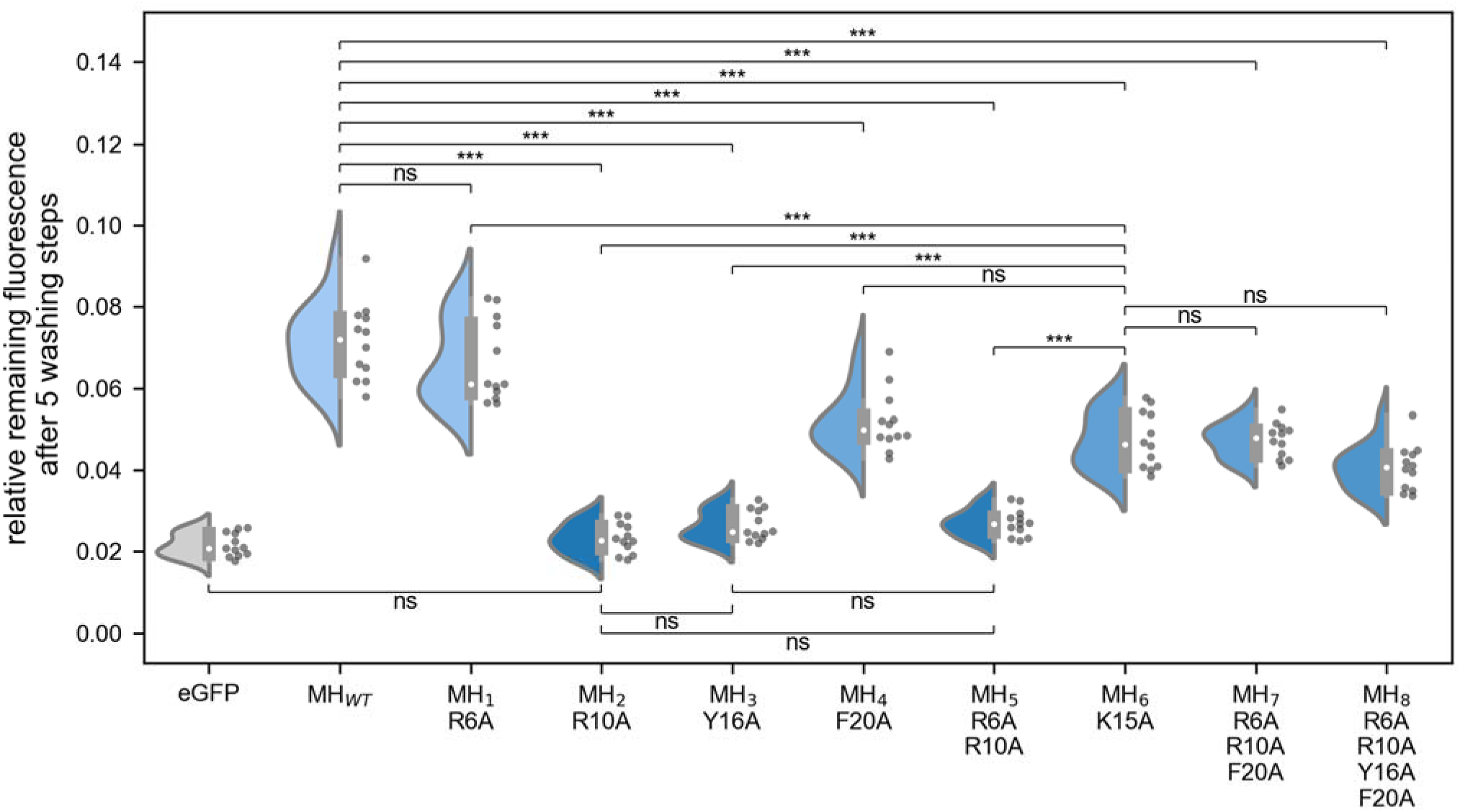
Binding of eGFP, eGFP-MacHis wild-type (MH_WT_), and eGFP-MacHis variants (MH_1-8_) to extracted cuticular wax from apple leaves. Binding was quantified by measuring the remaining fluorescence of the fusion partner eGFP after five washing steps on apple leaf cuticular wax in 96-well-MTPs (4 wells, 3 replicates). The similarity of the obtained distributions is evaluated using a two-sided Kolmogorov-Smirnov test; corresponding *p*-values are provided above the horizontal lines (ns: “not significant”, ***: p ≤ 0.001).

Significant differences in binding strength with regard to the wild-type and the negative control MH_6_ were observed for variants MH_2_, MH_3_, and MH_5._ These variants showed the overall lowest binding strength with a decrease of 97 %, 90 %, and 89 % in relative fluorescence compared to the wild-type, respectively, making them indistinguishable from pure eGFP. The variant MH_4_ as well as the negative control MH_6_ and the variants with multiple substitutions (MH_7_ and MH_8_) show a significant decrease in binding compared to the wild-type, however, to a lesser extent than variants MH_2_, MH_3_, and MH_5_. No significant difference in wax binding compared to the wild-type, however, was observed for variant MH_1_.

While single substitutions can lead to up to > 90 % decrease in binding, substitutions of several residues, including the single most impactful substitutions on positions 10 and 16, led to a decrease of at most 61 % (MH_7, 8_) except for MH_5_ (decrease of 89 %) compared to the wild-type. However, MH_7_ and MH_8_ do not show a significant difference in comparison to the negative control MH_6_. For MH_5_, the mutation R6A hardly influences the loss in binding strength caused by the mutation R10A, as expected from the behavior of MH_1_, leading to a significant decrease in binding compared to both the wild-type and the negative control. By contrast, in variants MH_7_ and MH_8_, the additional substitutions unexpectedly counterbalance the decrease in binding strength caused by the single substitutions at positions 10 and 16.

Out of the predicted preferred residues, identified from formed contacts, the two in the center (R10, Y16) show a significant effect on the binding strength, but the two on the sides (R6, F20) do not. This suggests that the contacts formed by the latter originate as a secondary effect from the contacts formed by R10 and Y16 and the rigid secondary structure of the α-helix. The finding is reminiscent of the O-ring hypothesis in protein-protein interfaces^62^ and indicates that more detailed energetic evaluations of residue contributions to binding^63^ may be necessary to evaluate AP-wax binding. Still, the computational prediction of binding-relevant residues provides a 117 times more likely identification of the two key residues (R10 and Y16) out of four possible suggestions than a random drawing, considering that the chance for finding the identified two key residues (assuming these are the only residues of major importance for binding) by randomly choosing four out of 38 possible mutation sides amounts to *h*(2|38;2;4) = 0.85 % based on the hypergeometric distribution. The two key residues are found in 53.8 % of the time when randomly generating 28 variants (*h*(2|38;2;28) = 53.8 %) and, therefore, the prediction using the computational models reduces the experimental burden by a factor of ∼7 (28 random variants / 4 predicted variants) on average. As to the unexpectedly small effects found for MH_7_ and MH_8_, we can only speculate that the multiple substitutions may lead to conformational changes of the AP and a differential binding mode, which was not captured in the MD simulations of the wild-type AP, suggesting that MD simulations of the variant APs are needed in such cases. In addition, in MH_1_, the mutation is close to the end connected to the stiff helical spacer, which in turn is connected to the eGFP, potentially restricting the N-terminal end of MacHis to form contacts with the cuticular wax.

### Binding Strengths Estimated via Adaptive Steered MD Match Experimentally Determined Binding Strengths of Aps

To estimate the individual binding properties of the APs, we performed adaptive-steered MD simulations. By applying a pseudo-force, we simulated the desorption of the AP from the leaf wax (Figure 7). The nonequilibrium work performed on the system can be related to the free energy difference between the bound and free states using Jarzynski’s equality (eq. 1, Figure 7 (inlay, blue transparent lines), Figure 8A). From the work calculated for each trajectory within each stage, we can deduce the PMF(*d*) for each stage as a function of the distance *d* of the AP’s geometric center from the surface of the leaf model (eq. 6, Figure 7 (inlay, black solid line), Figure 8A).

**Figure 7.**
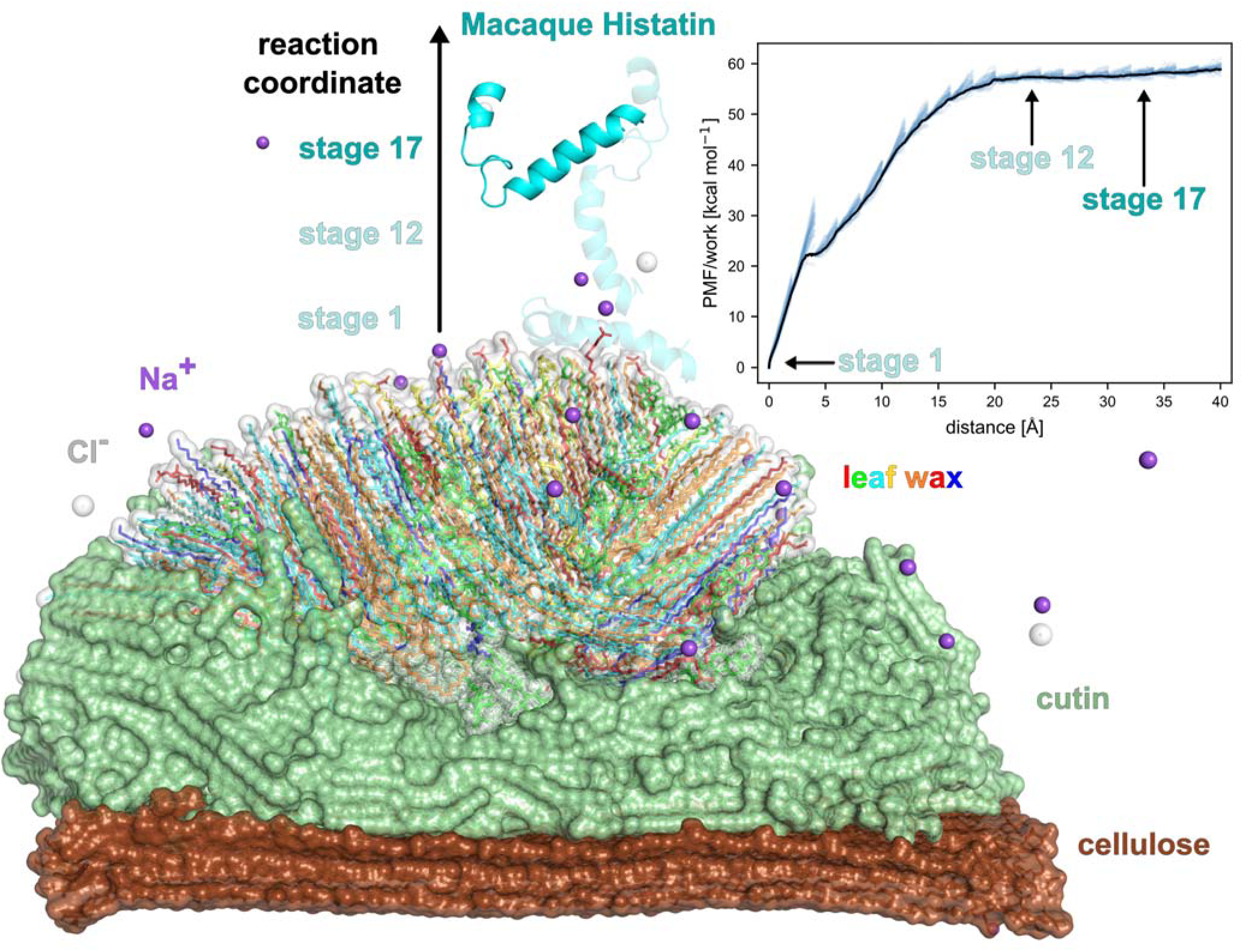
Adaptive steered MD of the desorption process of MacHis (cyan) from the leaf surface model. The potential of mean force (PMF, black solid line) of desorption determined as the average of the work of each stage (blue transparent lines) is depicted as inlay. The starting structure (stage 1) of MacHis as well as a structure selected from stage 12 of the steered MD are shown translucent, the completely detached AP is shown opaque (stage 17); corresponding stages are highlighted in the PMF profile.

**Figure 8.**
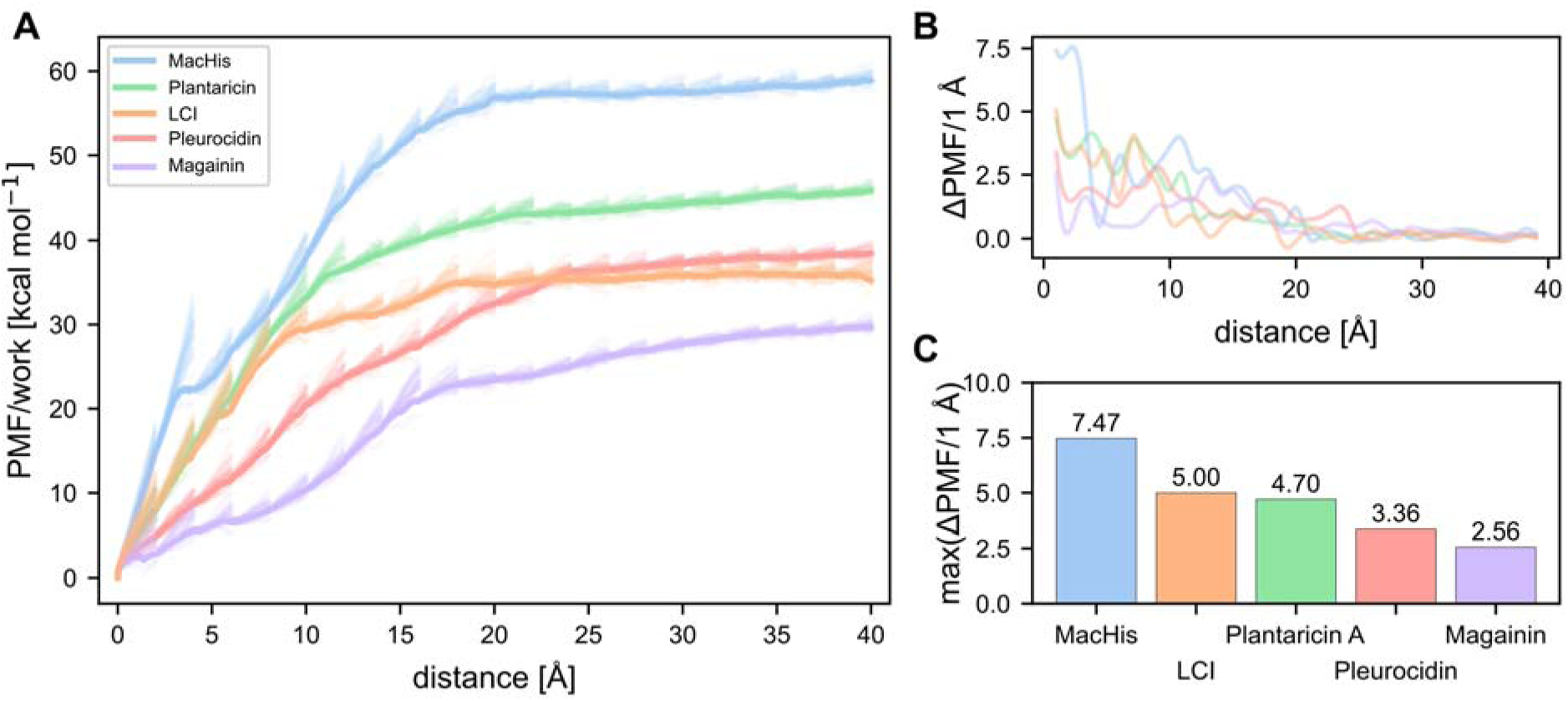
**A)** Potential of mean force (PMF) profiles for the desorption of MacHis (blue), Plantaricin A (green), LCI (orange), Pleurocidin (red), and Magainin (purple) as determined by adaptive steered MDs. The work of the 25 individual replicas per stage are depicted as transparent, and the PMF is shown opaque. **B)** Force profile as difference quotient ΔPMF/1 Å, where the denominator relates to the distance difference with respect to the leaf surface. **C)** Bar plot depicting the maxima of the forces shown in panel **B**.

The higher the work needed to transfer the AP from the wax surface into solution, the better the adhesion of the AP to the leaf surface. By determining the derivative of the applied work with respect to the distance, we obtain the force profile of the desorption process (Figure 8B), which can be used to determine a potential force threshold needed to release the AP from the leaf wax (Figure 8C). We performed the adaptive steered MDs for five APs, covering the range from weak to strong binding (Magainin, Pleurocidin, Plantaricin A, MacHis, and LCI).

The ASMD simulations reveal distinct differences in the PMFs for releasing the APs from the wax surface. The forced desorption of MacHis requires the most work (∼60 kcal mol^--1^), followed by Plantaricin A (∼40 kcal mol^--1^), LCI and Pleurocidin (∼35 kcal mol^--1^ each), and Magainin (∼30 kcal mol^--1^ each) (Figure 8A). As to the maximal force required during desorption (Figure 8B), MacHis and Magainin show the largest and smallest values, in line with the required work. The order of the other three APs is LCI > Plantaricin A > Pleurocidin, with the first two showing very similar values. The force needed to move the AP from the leaf surface is indicative of the steepness of the potential energy barriers along the reaction coordinate.

To assess our model’s capability to predict and rank binding affinities for structurally different APs qualitatively, all five APs were analyzed for their binding performance in the MTP-based binding assay. For a more stable production of all contigs in *E. coli*, the TEV cleavage site was excised using non-continuous (loop-out) primers during a PCR. As done in the alanine scanning, the APs’ binding strength was characterized by quantifying eGFP fluorescence in apple leaf wax pre-coated 96-well-PP-MTPs using the assay described above. The binding of only eGFP and eGFP-APs corresponds to the determined relative fluorescence in each well (4 wells, 3 replicates) in relation to the applied amount of AP. The results of the MTP assay of all eGFP-APs are depicted in Figure 9A (raw data is provided in Tables S7-S9).

**Figure 9.**
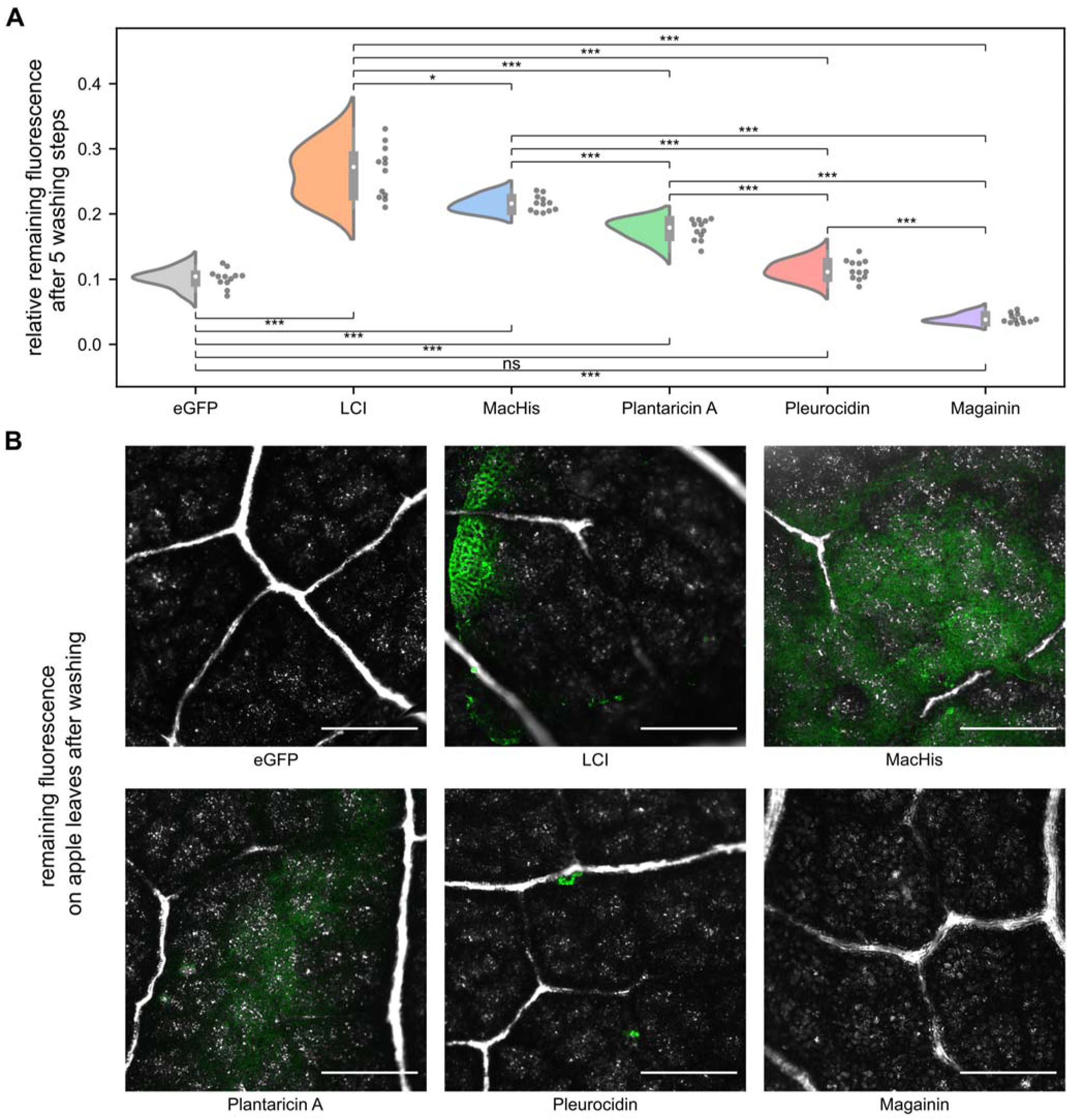
**A)** Binding of eGFP-APs to extracted cuticular wax from apple leaves. AP binding was quantified by measuring the remaining fluorescence of the reporter protein eGFP after five washing steps on apple leaf cuticular wax in 96-well-MTPs (4 wells, 3 replicates). The similarity of the obtained distributions is evaluated using a two-sided Kolmogorov-Smirnov test; corresponding *p*-values are provided above the corresponding horizontal lines (ns: “not significant”, *: p ≤0.05, ***: p ≤ 0.001). **B)** Qualitative assessment of eGFP-AP binding on apple leaves after washing using confocal microscopy. Immobilization of eGFP-control and eGFP-APs to the surface of apple leaves was investigated by incubation (50 µL, 5 min, ambient temperature) followed by one washing step (1 mL, 50 mM Tris-HCl buffer pH 8.0). Scale bars represent 500 µm in all images.

After five washing steps, LCI and MacHis show the highest binding strength to surface apple leaf wax according to the relative remaining fluorescence, followed by Plantaricin A, Pleurocidin, and Magainin (Figure 9A); the relative remaining fluorescence of Pleurocidin is not significantly different from the pure eGFP, and the one of Magainin is significantly worse (results after two washing steps are depicted in Figure S6). These results compare very favorably to the computed maximal forces required during desorption in that the same order of binding strength is obtained except for LCI and MacHis (Figure 8C).

During the production of eGFP-Plantaricin A and eGFP-Magainin in *E. coli*, a degradation band was observed in the SDS-PAGE at a size corresponding to eGFP. The eGFP-AP concentration was normalized based on the concentration quantified by SDS-PAGE using ImageJ/FiJi (Figure S7). Despite the normalized eGFP-AP concentration, the different peptide samples contained a varying amount of eGFP (eGFP-Magainin: ∼75 %, eGFP-Plantaricin A: ∼10 %). eGFP could interfere with the binding of Magainin and Plantaricin A leading to a possibly lower binding strength.

The quantitative results from the MTP assay agree well with qualitative results obtained from fluorescence microscopy of apple leaves first treated with the eGFP-AP solutions and subsequently washed (Figure 9B), supporting that the wax-coated MTPs adequately mimic the waxy surface of an apple leaf.

Overall, the results from the MTP assay largely match the computational predictions and the qualitative observations on entire apple leaves. The good agreement with the (AS)MD simulations supports the validity of our atomistic leaf model and the computational approach. To further increase the precision of the PMFs obtained from the ASMD simulations, ideally, several PMFs obtained from different starting conformations using different pulling directions should be calculated. However, ASMD simulations are computationally costly for systems comprising more than 1,000,000 atoms.

## 5 Conclusion

In this work, we present a workflow for the selection of suitable APs and rational improvement of their binding properties towards specific plant leaves using a multidisciplinary approach. For this, we created the, to our knowledge, first multi-layered model of an apple leaf surface, which can be used to investigate interactions of agrochemicals or nutrients with leaf surfaces at an atomistic level. The model consists of three layers: cellulose, cutin, and a surface wax layer. All layers can be modified and adjusted to reflect the properties of different plant leaves. With the leaf wax composition of apple leaves determined with GC/FID and GC/MS, we could tailor our atomistic model towards that specific leaf type, enabling the realistic modeling of adsorption properties.

To validate our model, we focused on investigating the adhesion of APs to the cuticular wax layer of the model. APs are versatile and well-studied adhesion promotors that can be applied in a variety of fields, including novel plant protection technologies.^9^ For complex biological surfaces such as a leaf surface, however, the mechanisms of adsorption at the atomistic level have remained elusive. To elucidate the adsorption, we first used our atomistic leaf surface model to identify key residues of the AP MacHis that preferentially interact with the wax layer in MD simulations. The identified residues were further investigated experimentally using alanine scanning in combination with a newly developed MTP assay to rapidly quantify the surface wax binding properties of different APs or AP variants. With the obtained knowledge and the identified key binding residues of MacHis, we can tune binding strength to improve rainfastness to match application demands. For MacHis, we found that aromatic and positively charged amino acids on one side of the helix majorly contribute to the binding towards surface wax of apple leaves. Therefore, engineered anchor peptides might become promising alternatives for polymeric adhesion promoters and will pave the way to develop microplastics-free and biodegradable plant protection products.

Second, we probed our workflow for use in AP screening. For this, we performed ASMD simulations to determine the PMF of desorption of five different APs from the leaf surface, ranging from weak to strong binding APs. The results were validated with the novel MTP assay. The quantitative experimental results match well with the computational predictions as well as with qualitative results obtained from experiments using entire leaves.

The established workflow opens up avenues for further experimental and computational studies revolving around foliar applications, such as optimizing APs and investigating the adsorption, incorporation, or diffusion of herbicides, fungicides, or nutrients on, into, or through the surface wax, cutin, or cellulose. Finally, the generated atomistic models and the MTP assay can be adapted to accommodate different plant types and growing conditions.

## 6 Acknowledgements

We gratefully acknowledge the computational support provided by the “Center for Information and Media Technology” (ZIM) at the Heinrich Heine University Düsseldorf and the computing time provided by the John von Neumann Institute for Computing (NIC) on the supercomputer JUWELS at Jülich Supercomputing Centre (JSC) (user IDs: HKF7, VSK33, project “microgels”).

## ASSOCIATED CONTENT

### Supporting Information

Additional information on the surface wax composition, nucleotide and primer sequences, SDS-PAGE data, contacts of the investigated APs with the leaf, bootstrapping analyses on the identified binding promoting residues, stability of the MacHis binding positions during MDs, and the remaining fluorescence of the eGFP-AP constructs after two washing steps is provided in the SI.

This material is available free of charge via the internet at http://pubs.acs.org.

## AUTHOR INFORMATION

## AUTHOR CONTRIBUTIONS

JD generated the atomistic models, performed the computational studies, analyzed results, and wrote the manuscript; CB generated and produced the anchor peptides and performed binding experiments, analyzed results, and wrote the manuscript; LG performed the immobilization of eGFP-APs on apple leaves; JB performed wax extraction; SP supervised field experiments from where apple leaf samples and leaf wax data were made available and wrote part of the manuscript; VZ and LS performed the wax analysis and wrote part of the manuscript; FJ supervised experimental work, analyzed results and wrote parts of the manuscript; TK supervised experimental work and analyzed results; US supervised experimental work, analyzed results, and wrote parts of the manuscript; HG conceived the study, supervised and managed the project, and wrote the manuscript. All authors have given approval to the final version of the manuscript.

## NOTES

The authors declare no competing financial interest.

## 8 TOC Figure

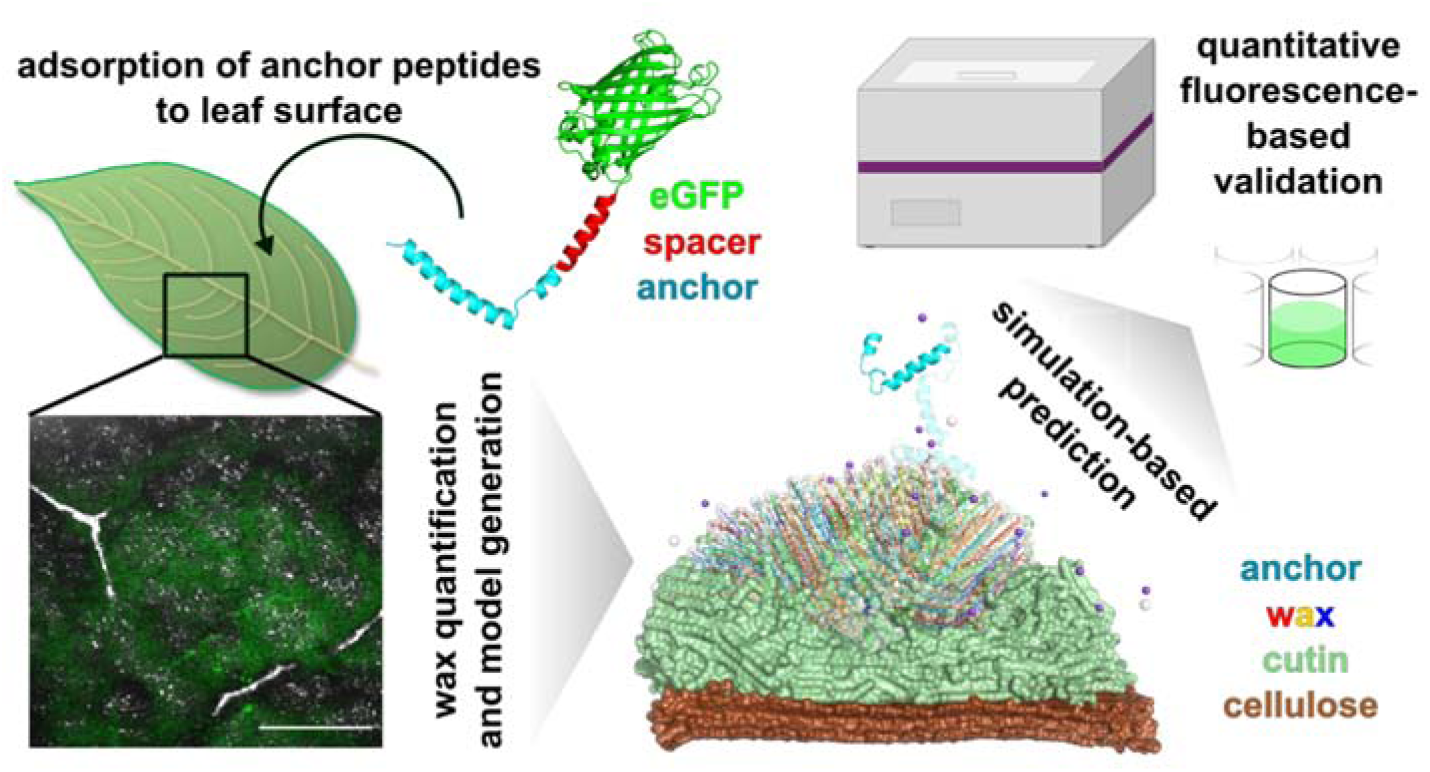

**For Table of Contents Only**

## References

(1) Cakmak, I., Enrichment of cereal grains with zinc: Agronomic or genetic biofortification? Plant Soil 2008, 302 (1), 1–17.

(2) Frank, C. R.; Loston, R.; Donald, P.; Len, P.; Richard, B., Increasing Postemergence Herbicide Efficacy and Rainfastness with Silicone Adjuvants. Weed Technol. 1990, 4 (3), 576–580.

(3) Hunsche, M.; Bringe, K.; Schmitz-Eiberger, M.; Noga, G., Leaf surface characteristics of apple seedlings, bean seedlings and kohlrabi plants and their impact on the retention and rainfastness of mancozeb. Pest Manage. Sci. 2006, 62 (9), 839–847.

(4) Vicent, A.; Armengol, J.; García-Jiménez, J., Rain Fastness and Persistence of Fungicides for Control of Alternaria Brown Spot of Citrus. Plant Dis. 2007, 91 (4), 393–399.

(5) Symonds, B. L.; Thomson, N. R.; Lindsay, C. I.; Khutoryanskiy, V. V., Rainfastness of Poly(vinyl alcohol) Deposits on Vicia faba Leaf Surfaces: From Laboratory-Scale Washing to Simulated Rain. ACS Appl. Mater. Interfaces 2016, 8 (22), 14220–14230.

(6) Gao, Y.; Li, X.; He, L.; Li, B.; Mu, W.; Liu, F., Role of Adjuvants in the Management of Anthracnose—Change in the Crystal Morphology and Wetting Properties of Fungicides. J. Agric. Food Chem. 2019, 67 (33), 9232–9240.

(7) Krishna, N. R.; Singh, M., Organosilicone Adjuvant Effects on Glyphosate Efficacy and Rainfastness. Weed Technol. 1992, 6 (2), 361–365.

(8) Symonds, Brett L.; Lindsay, C. I.; Thomson, N. R.; Khutoryanskiy, V. V., Chitosan as a rainfastness adjuvant for agrochemicals. RSC Adv. 2016, 6 (104), 102206–102213.

(9) Meurer, R. A.; Kemper, S.; Knopp, S.; Eichert, T.; Jakob, F.; Goldbach, H. E.; Schwaneberg, U.; Pich, A., Biofunctional Microgel-Based Fertilizers for Controlled Foliar Delivery of Nutrients to Plants. Angew. Chem. Int. Ed. 2017, 56 (26), 7380–7386.

(10) Schwinges, P.; Pariyar, S.; Jakob, F.; Rahimi, M.; Apitius, L.; Hunsche, M.; Schmitt, L.; Noga, G.; Langenbach, C.; Schwaneberg, U.; Conrath, U., A bifunctional dermaseptin–thanatin dipeptide functionalizes the crop surface for sustainable pest management. Green Chem. 2019, 21 (9), 2316–2325.

(11) Rübsam, K.; Weber, L.; Jakob, F.; Schwaneberg, U., Directed evolution of polypropylene and polystyrene binding peptides. Biotechnol. Bioeng. 2018, 115 (2), 321–330.

(12) Rübsam, K.; Davari, M. D.; Jakob, F.; Schwaneberg, U., KnowVolution of the polymer-binding peptide LCI for improved polypropylene binding. Polymers 2018, 10 (4), 423.

(13) Rübsam, K.; Stomps, B.; Böker, A.; Jakob, F.; Schwaneberg, U., Anchor peptides: A green and versatile method for polypropylene functionalization. Polymer 2017, 116, 124–132.

(14) Nöth, M.; Hussmann, L.; Belthle, T.; El-Awaad, I.; Davari, M. D.; Jakob, F.; Pich, A.; Schwaneberg, U., MicroGelzymes: pH-Independent Immobilization of Cytochrome P450 BM3 in Microgels. Biomacromolecules 2020, 21 (12), 5128–5138.

(15) Dedisch, S.; Wiens, A.; Davari, M. D.; Söder, D.; Rodriguez-Emmenegger, C.; Jakob, F.; Schwaneberg, U., Matter-tag: A universal immobilization platform for enzymes on polymers, metals, and silicon-based materials. Biotechnol. Bioeng. 2020, 117 (1), 49–61.

(16) Apitius, L.; Buschmann, S.; Bergs, C.; Schönauer, D.; Jakob, F.; Pich, A.; Schwaneberg, U., Biadhesive Peptides for Assembling Stainless Steel and Compound Loaded Micro-Containers. Macromol. Biosci. 2019, 19 (9), 1900125.

(17) Büscher, N.; Sayoga, G. V.; Rübsam, K.; Jakob, F.; Schwaneberg, U.; Kara, S.; Liese, A., Biocatalyst Immobilization by Anchor Peptides on an Additively Manufacturable Material. Org. Process Res. Dev. 2019, 23 (9), 1852–1859.

(18) Grimm, A. R.; Sauer, D. F.; Mirzaei Garakani, T.; Rübsam, K.; Polen, T.; Davari, M. D.; Jakob, F.; Schiffels, J.; Okuda, J.; Schwaneberg, U., Anchor Peptide-Mediated Surface Immobilization of a Grubbs-Hoveyda-Type Catalyst for Ring-Opening Metathesis Polymerization. Bioconj. Chem. 2019, 30 (3), 714–720.

(19) Schwemmer, T.; Baumgartner, J.; Faivre, D.; Börner, H. G., Peptide-Mediated Nanoengineering of Inorganic Particle Surfaces: A General Route toward Surface Functionalization via Peptide Adhesion Domains. J. Am. Chem. Soc. 2012, 134 (4), 2385–2391.

(20) Gao, X.; Ran, N.; Dong, X.; Zuo, B.; Yang, R.; Zhou, Q.; Moulton, H. M.; Seow, Y.; Yin, H., Anchor peptide captures, targets, and loads exosomes of diverse origins for diagnostics and therapy. Sci. Transl. Med. 2018, 10 (444), eaat0195.

(21) Gong, W.; Wang, J.; Chen, Z.; Xia, B.; Lu, G., Solution Structure of LCI, a Novel Antimicrobial Peptide from Bacillus subtilis. Biochemistry 2011, 50 (18), 3621–3627.

(22) Hess, F. D.; Falk, R. H., Herbicide Deposition on Leaf Surfaces. Weed Sci. 1990, 38 (3), 280–288.

(23) Kolattukudy, P. E., Biopolyester Membranes of Plants: Cutin and Suberin. Science 1980, 208 (4447), 990–1000.

(24) Bakan, B.; Marion, D., Assembly of the cutin polyester: from cells to extracellular cell walls. Plants 2017, 6 (4), 57.

(25) Gomes, T. C. F.; Skaf, M. S., Cellulose-Builder: A toolkit for building crystalline structures of cellulose. J. Comput. Chem. 2012, 33 (14), 1338–1346.

(26) Kirschner, K. N.; Yongye, A. B.; Tschampel, S. M.; González-Outeiriño, J.; Daniels, C. R.; Foley, B. L.; Woods, R. J., GLYCAM06: A generalizable biomolecular force field. Carbohydrates. J. Comput. Chem. 2008, 29 (4), 622–655.

(27) Fich, E. A.; Segerson, N. A.; Rose, J. K. C., The Plant Polyester Cutin: Biosynthesis, Structure, and Biological Roles. Annu. Rev. Plant Biol. 2016, 67 (1), 207–233.

(28) Schrödinger Maestro, Schrödinger: New York, NY, 2019.

(29) Martínez, L.; Andrade, R.; Birgin, E. G.; Martínez, J. M., PACKMOL: A package for building initial configurations for molecular dynamics simulations. J. Comput. Chem. 2009, 30 (13), 2157–2164.

(30) Jorgensen, W. L.; Chandrasekhar, J.; Madura, J. D.; Impey, R. W.; Klein, M. L., Comparison of simple potential functions for simulating liquid water. J. Chem. Phys. 1983, 79 (2), 926–935.

(31) Jakalian, A.; Bush, B. L.; Jack, D. B.; Bayly, C. I., Fast, efficient generation of high-quality atomic charges. AM1-BCC model: I. Method. J. Comput. Chem. 2000, 21 (2), 132–146.

(32) Jakalian, A.; Jack, D. B.; Bayly, C. I., Fast, efficient generation of high-quality atomic charges. AM1-BCC model: II. Parameterization and validation. J. Comput. Chem. 2002, 23 (16), 1623–1641.

(33) Wang, J.; Wang, W.; Kollman, P. A.; Case, D. A., Automatic atom type and bond type perception in molecular mechanical calculations. J. Mol. Graph. Model. 2006, 25 (2), 247–260.

(34) Wang, J.; Wolf, R. M.; Caldwell, J. W.; Kollman, P. A.; Case, D. A., Development and testing of a general amber force field. J. Comput. Chem. 2004, 25 (9), 1157–1174.

(35) Berman, H. M.; Westbrook, J.; Feng, Z.; Gilliland, G.; Bhat, T. N.; Weissig, H.; Shindyalov, I. N.; Bourne, P. E., The Protein Data Bank. Nucleic Acids Res. 2000, 28 (1), 235–242.

(36) Mulnaes, D.; Porta, N.; Clemens, R.; Apanasenko, I.; Reiners, J.; Gremer, L.; Neudecker, P.; Smits, S. H. J.; Gohlke, H., TopModel: Template-Based Protein Structure Prediction at Low Sequence Identity Using Top-Down Consensus and Deep Neural Networks. J. Chem. Theory Comput. 2020, 16 (3), 1953–1967.

(37) Case, D. A.; Cheatham III, T. E.; Darden, T.; Gohlke, H.; Luo, R.; Merz Jr., K. M.; Onufriev, A.; Simmerling, C.; Wang, B.; Woods, R. J., The Amber biomolecular simulation programs. J. Comput. Chem. 2005, 26 (16), 1668–1688.

(38) Case, D. A.; Ben-Shalom, I. Y.; Brozell, S. R.; Cerutti, D. S.; Cheatham III, T. E.; Cruzeiro, V. W. D.; Darden, T. A.; Duke, R. E.; Ghoreishi, D.; Gilson, M. K.; Gohlke, H.; Goetz, A. W.; Greene, D.; Harris, R.; Homeyer, N.; Izadi, S.; Kovalenko, A.; Kurtzman, T.; Lee, T. S.; LeGrand, S.; Li, P.; Lin, C.; Liu, J.; Luchko, T.; Luo, R.; Mermelstein, D. J.; Merz, K. M.; Miao, Y.; Monard, G.; Nguyen, C.; Nguyen, H.; Omelyan, I.; Onufriev, A.; Pan, F.; Qi, R.; Roe, D. R.; Roitberg, A.; Sagui, C.; Schott-Verdugo, S.; Shen, J.; Simmerling, C. L.; Smith, J.; Salomon-Ferrer, R.; Swails, J.; Walker, R. C.; Wang, J.; Wei, H.; Wolf, R. M.; Wu, X.; Xiao, L.; York, D. M.; Kollman, P. A. AMBER, University of California: San Francisco, 2021.

(39) Salomon-Ferrer, R.; Götz, A. W.; Poole, D.; Le Grand, S.; Walker, R. C., Routine Microsecond Molecular Dynamics Simulations with AMBER on GPUs. 2. Explicit Solvent Particle Mesh Ewald. J. Chem. Theory Comput. 2013, 9 (9), 3878–3888.

(40) Le Grand, S.; Götz, A. W.; Walker, R. C., SPFP: Speed without compromise—A mixed precision model for GPU accelerated molecular dynamics simulations. Comput. Phys. Commun. 2013, 184 (2), 374–380.

(41) Maier, J. A.; Martinez, C.; Kasavajhala, K.; Wickstrom, L.; Hauser, K. E.; Simmerling, C., ff14SB: Improving the Accuracy of Protein Side Chain and Backbone Parameters from ff99SB. J. Chem. Theory Comput. 2015, 11 (8), 3696–3713.

(42) Darden, T.; York, D.; Pedersen, L., Particle mesh Ewald: An N·log(N) method for Ewald sums in large systems. J. Chem. Phys. 1993, 98 (12), 10089–10092.

(43) Ryckaert, J.-P.; Ciccotti, G.; Berendsen, H. J. C., Numerical integration of the cartesian equations of motion of a system with constraints: molecular dynamics of n-alkanes. J. Comput. Phys. 1977, 23 (3), 327–341.

(44) Berendsen, H. J. C.; Postma, J. P. M.; Gunsteren, W. F. v.; DiNola, A.; Haak, J. R., Molecular dynamics with coupling to an external bath. J. Chem. Phys. 1984, 81 (8), 3684–3690.

(45) Ozer, G.; Valeev, E. F.; Quirk, S.; Hernandez, R., Adaptive Steered Molecular Dynamics of the Long-Distance Unfolding of Neuropeptide Y. J. Chem. Theory Comput. 2010, 6 (10), 3026–3038.

(46) Ozer, G.; Quirk, S.; Hernandez, R., Adaptive steered molecular dynamics: Validation of the selection criterion and benchmarking energetics in vacuum. J. Chem. Phys. 2012, 136 (21), 215104.

(47) Jarzynski, C., Nonequilibrium equality for free energy differences. Phys. Rev. Lett. 1997, 78 (14), 2690.

(48) Park, S.; Schulten, K., Calculating potentials of mean force from steered molecular dynamics simulations. J. Chem. Phys. 2004, 120 (13), 5946–5961.

(49) Ozer, G.; Quirk, S.; Hernandez, R., Thermodynamics of Decaalanine Stretching in Water Obtained by Adaptive Steered Molecular Dynamics Simulations. J. Chem. Theory Comput. 2012, 8 (11), 4837–4844.

(50) Ahmad, S.; Strunk, C. H.; Schott-Verdugo, S. N.; Jaeger, K.-E.; Kovacic, F.; Gohlke, H., Substrate Access Mechanism in a Novel Membrane-Bound Phospholipase A of Pseudomonas aeruginosa Concordant with Specificity and Regioselectivity. J. Chem. Inf. Model. 2021, 61 (11), 5626–5643.

(51) Roe, D. R.; Cheatham, T. E., PTRAJ and CPPTRAJ: Software for Processing and Analysis of Molecular Dynamics Trajectory Data. J. Chem. Theory Comput. 2013, 9 (7), 3084–3095.

(52) Arai, R.; Ueda, H.; Kitayama, A.; Kamiya, N.; Nagamune, T., Design of the linkers which effectively separate domains of a bifunctional fusion protein. Protein Engineering, Design and Selection 2001, 14 (8), 529–532.

(53) Kapust, R. B.; Tözsér, J.; Fox, J. D.; Anderson, D. E.; Cherry, S.; Copeland, T. D.; Waugh, D. S., Tobacco etch virus protease: mechanism of autolysis and rational design of stable mutants with wild-type catalytic proficiency. Protein Engineering, Design and Selection 2001, 14 (12), 993–1000.

(54) Blanusa, M.; Schenk, A.; Sadeghi, H.; Marienhagen, J.; Schwaneberg, U., Phosphorothioate-based ligase-independent gene cloning (PLICing): An enzyme-free and sequence-independent cloning method. Anal. Biochem. 2010, 406 (2), 141–146.

(55) Bringe, K.; Schumacher, C. F. A.; Schmitz-Eiberger, M.; Steiner, U.; Oerke, E.-C., Ontogenetic variation in chemical and physical characteristics of adaxial apple leaf surfaces. Phytochemistry 2006, 67 (2), 161–170.

(56) Sterling, T. M., Mechanisms of Herbicide Absorption across Plant Membranes and Accumulation in Plant Cells. Weed Sci. 1994, 42 (2), 263–276.

(57) Mishra, V. K.; Upadhyay, A. R.; Tripathi, B. D., Bioaccumulation of heavy metals and two organochlorine pesticides (DDT and BHC) in crops irrigated with secondary treated waste water. Environ. Monit. Assess. 2009, 156 (1), 99–107.

(58) Liu, Q.; Liu, Y.; Dong, F.; Sallach, J. B.; Wu, X.; Liu, X.; Xu, J.; Zheng, Y.; Li, Y., Uptake kinetics and accumulation of pesticides in wheat (Triticum aestivum L.): Impact of chemical and plant properties. Environ. Pollut. 2021, 275, 116637.

(59) Izvekov, S.; Voth, G. A., A Multiscale Coarse-Graining Method for Biomolecular Systems. J. Phys. Chem. B 2005, 109 (7), 2469–2473.

(60) Saunders, M. G.; Voth, G. A., Coarse-Graining Methods for Computational Biology. Annual Review of Biophysics 2013, 42 (1), 73–93.

(61) Kabsch, W.; Sander, C., Dictionary of protein secondary structure: Pattern recognition of hydrogen-bonded and geometrical features. Biopolymers 1983, 22 (12), 2577–2637.

(62) Li, Y.; Huang, Y.; Swaminathan, C. P.; Smith-Gill, S. J.; Mariuzza, R. A., Magnitude of the Hydrophobic Effect at Central versus Peripheral Sites in Protein-Protein Interfaces. Structure 2005, 13 (2), 297–307.

(63) Gohlke, H.; Kiel, C.; Case, D. A., Insights into Protein–Protein Binding by Binding Free Energy Calculation and Free Energy Decomposition for the Ras–Raf and Ras–RalGDS Complexes. J. Mol. Biol. 2003, 330 (4), 891–913.

