## Supplemental Information for "Rational design yields molecular insights on leaf binding of the anchor peptide Macaque Histatin"

### Table of Contents

**Supplementary Tables** ........ **5**

**Table S1.** Composition of apple (*Malus domestica*; cultivar 'Bittenfelder') cuticular wax obtained via GC/FID and GC/MS. ........ 5

**Table S2.** Nucleotide sequences of ordered synthetic genes of LCI (UniProt ID: P82243), Macaque Histatin (MacHis; UniProt ID: P34084), Magainin 2 (UniProt ID: P11006), Plantaricin A (PlnA; UniProt ID: P80214), and Pleurocidin (UniProt ID: P81941). Underlined sequence encodes for the 17 Helix spacer, grey nucleotides encodes for the TEV protease cleavage site, bold nucleotides represent the peptide sequences. The genes were codon-optimized for *E. coli* and synthesized by GeneScript (Nanjing, China). ........ 6

**Figure S1.** Residue-wise relative contacts of (A) LCI, (B) Macaque Histatin, (C) Magainin, (D) Pleurocidin, and (E) Plantaricin with the wax molecules within 10 x 250 ns long MD simulations of anchor peptide adsorption. Error bars denote the SEM. Geometrical analyses of the MD simulations were performed using cpptraj.[1] On top of the plots, the secondary structure of the APs is indicated (wavy line: α-helix, arrow: β-strand) as determined by DSSP[2] analysis of the protein structures. ........................................................................................ 11

**Figure S2.** Bootstrapping analysis for identifying residues important for cuticular wax binding of MacHis in dependence of the sample size. Samples of different sizes (5, 10, 15, 20, 25, 29) were drawn 10,000 times with replacement from the original set containing 29 samples. For each sample set size, the overlap with the top three (**A**, **B**) and top five (**C**, **D**) identified residues using the original set is calculated. The probability of identifying exactly X residues out of the three/five is given in **A** and **C**, the probability of identifying at least X residues out of the three/five is given in **B** and **D** for each sample size. ....................................... 12

**Figure S3.** Bootstrapping analysis for identifying residues important for cuticular wax binding of LCI in dependence of the sample size. Samples of different sizes (5, 15, 25, 50, 100, 200, 281) were drawn 10,000 times with replacement from the original set containing 281 samples. For each sample set size, the overlap with the top three (**A**, **B**) and top five (**C**, **D**) identified residues using the original set is calculated. The probability of identifying exactly X residues out of the three/five is given in **A** and **C**, the probability of identifying at least X residues out of the three/five is given in **B** and **D** for each sample size. ....................... 13

### Supplementary Tables

**Table S1.** Composition of apple (*Malus domestica*; cultivar 'Bittenfelder') cuticular wax obtained via GC/FID and GC/MS.

| component | | amount[a] replica 1 | 2 | 3 | 4 | 5 | mean[a] | SEM[a,b] |
|---|---|---|---|---|---|---|---|---|
| fatty acid | $C_{16}$ | 0.120 | 0.130 | 0.108 | 0.187 | 0.161 | 0.141 | 0.014 |
| | $C_{18}$ | 0.132 | 0.131 | 0.108 | 0.226 | 0.180 | 0.155 | 0.021 |
| | $C_{20}$ | 0.016 | 0.016 | 0.018 | 0.021 | 0.074 | 0.029 | 0.011 |
| | $C_{22}$ | 0.012 | 0.028 | 0.023 | 0.029 | 0.050 | 0.028 | 0.006 |
| | $C_{24}$ | 0.023 | 0.040 | 0.064 | 0.026 | 0.035 | 0.037 | 0.007 |
| | $C_{26}$ | 0.095 | 0.129 | 0.152 | 0.105 | 0.110 | 0.118 | 0.010 |
| | $C_{28}$ | 0.083 | 0.153 | 0.102 | 0.103 | 0.081 | 0.105 | 0.013 |
| | $C_{30}$ | 0.095 | 0.068 | 0.038 | 0.031 | 0.053 | 0.057 | 0.011 |
| | $C_{32}$ | 0.015 | 0.042 | 0.048 | 0.060 | 0.031 | 0.039 | 0.008 |
| | $C_{34}$ | 0.029 | 0.055 | 0.044 | 0.044 | 0.044 | 0.043 | 0.004 |
| alcohol | $C_{26}$ | 0.574 | 0.510 | 0.812 | 0.534 | 0.664 | 0.619 | 0.055 |
| | $C_{27}$ | 0.018 | 0.023 | 0.033 | 0.024 | 0.024 | 0.024 | 0.002 |
| | $C_{28}$ | 0.281 | 0.248 | 0.390 | 0.260 | 0.352 | 0.306 | 0.028 |
| | $C_{29}$ | 0.028 | 0.006 | 0.056 | 0.065 | 0.046 | 0.040 | 0.010 |
| | $C_{30}$ | 0.245 | 0.238 | 0.481 | 0.282 | 0.392 | 0.327 | 0.047 |
| | $C_{31}$ | 0.021 | 0.129 | 0.067 | 0.104 | 0.016 | 0.067 | 0.022 |
| | $C_{32}$ | 0.338 | 0.517 | 0.823 | 0.559 | 0.652 | 0.578 | 0.080 |
| | $C_{33}$ | 0.004 | 0.057 | 0.029 | 0.026 | 0.015 | 0.026 | 0.009 |
| | $C_{34}$ | 0.073 | 0.109 | 0.255 | 0.110 | 0.108 | 0.131 | 0.032 |
| alkane | $C_{31}$ | 1.677 | 1.522 | 1.749 | 1.374 | 1.265 | 1.517 | 0.090 |
| | $C_{33}$ | 0.406 | 0.361 | 0.340 | 0.282 | 0.213 | 0.320 | 0.033 |
| sterol | campesterol | 0.227 | 0.160 | 0.104 | 0.203 | 0.108 | 0.161 | 0.025 |
| | beta-sitosterol | 0.643 | 0.341 | 0.872 | 0.769 | 0.657 | 0.656 | 0.089 |
| | stigmasterol | 0.093 | 0.005 | 0.102 | 0.157 | 0.055 | 0.083 | 0.025 |
| terpenoid | uvaol | 0.056 | 0.133 | 0.104 | 0.112 | 0.052 | 0.091 | 0.016 |
| | oleanolic acid | 0.034 | 0.116 | 0.066 | 0.085 | 0.071 | 0.074 | 0.013 |
| | ursolic acid | 0.069 | 0.063 | 0.198 | 0.068 | 0.107 | 0.101 | 0.026 |
| ketone | $C_{34}$ | 0.153 | 0.082 | 0.281 | 0.115 | 0.224 | 0.171 | 0.036 |
| | $C_{35}$ | 0.421 | 0.324 | 0.261 | 0.282 | 0.280 | 0.314 | 0.029 |
| ester | $C_{40}$ | 0.027 | 0.058 | 0.128 | 0.063 | 0.070 | 0.069 | 0.017 |
| | $C_{42}$ | 0.052 | 0.118 | 0.215 | 0.073 | 0.075 | 0.107 | 0.029 |
| | $C_{44}$ | 0.136 | 0.152 | 0.254 | 0.122 | 0.141 | 0.161 | 0.024 |
| | $C_{46}$ | 0.196 | 0.181 | 0.363 | 0.142 | 0.217 | 0.220 | 0.038 |
| | $C_{48}$ | 0.133 | 0.097 | 0.288 | 0.082 | 0.133 | 0.147 | 0.037 |

[a] In µg $cm^{-2}$.

[b] Standard error of the mean.

**Table S2.** Nucleotide sequences of ordered synthetic genes of LCI (UniProt ID: P82243), Macaque Histatin (MacHis; UniProt ID: P34084), Magainin 2 (UniProt ID: P11006), Plantaricin A (PlnA; UniProt ID: P80214), and Pleurocidin (UniProt ID: P81941). The underlined sequence encodes for the 17 Helix spacer, grey nucleotides encode for the TEV protease cleavage site, bold nucleotides represent the peptide sequences. The genes were codon-optimized for *E. coli* and synthesized by GeneScript (Nanjing, China).

| |
|---|
| **LCI**<br>5'-<br>GCAGAAGCAGCAGCAAAAGAAGCCGCTGCCAAAGAAGCGGCAGCGAAAGCAGAAAATCTGTAT<br>TTTCAGGGT**GCCATTAAACTGGTTCAGAGCCCGAATGGTAATTTTGCAGCAAGCTTTGTTCT**<br>**GGATGGCACCAAATGGATCTTCAAAAGCAAATACTATGACAGCAGCAAAGGTTATTGGGTG**<br>**GGTATTTATGAAGTGTGGGATCGCAAA**TAATAA-3' |
| **Macaque Histatin**<br>5'-<br>GCAGAAGCAGCTGCCAAAGAAGCGGCAGCGAAAGAAGCGGCGGCCAAAGCCGAGAATCTGTAC<br>TTTCAGGGC**GATTCTCACGAAGAACGCCATCATGGTCGTCATGGTCACCACAAGTATGGCC**<br>**GCAAATTCCACGAGAAACATCACAGTCATCGTGGCTATCGCTCGAACTATCTGTACGACAA**<br>**C**TGATAA-3' |
| **Plantaricin A**<br>5'-<br>GCAGAAGCAGCAGCCAAAGAAGCTGCGGCGAAAGAAGCGGCAGCCAAAGCGGAGAACCTGTA<br>CTTTCAAGGC**AAAAGCAGTGCGTATTCCTTGCAAATGGGTGCCACCGCCATTAAACAGGTT**<br>**AAGAAACTGTTCAAGAAATGGGGCTGG**TAA-3' |
| **Pleurocidin**<br>5'-<br>GCAGAAGCAGCGGCAAAAGAGGCTGCTGCGAAAGAAGCTGCGGCCAAAGCGGAGAACCTGTAC<br>TTTCAGGGT**GGATGGGGCTCGTTCTTTAAGAAGGCTGCACATGTGGGCAAACACGTTGGGA**<br>**AAGCCGCCTTAACGCACTATCTG**TGATAA-3' |
| **Magainin**<br>5'-<br>GCAGAAGCAGCGGCCAAAGAAGCGGCAGCGAAAGAGGCTGCAGCGAAAGCCGAAAACCTGTAT<br>TTTCAGGGT**GGTATTGGGAAATTCCTGCATTCCGCGAAGAAATTCGGCAAAGCCTTTGTTG**<br>**GCGAGATCATGAACTCG**TGATAA-3' |

**Table S3.** Primer sequences for TEV site removal in a PCR and amino acid substitution into MacHis.

| Name | Sequence 5’-3’ |
|---|---|
| LCI-TEV-fw | GCCAAAGAAGCGGCAGCGAAAGCAGCCATTAAACTGGTTCAGAGCCCG |
| LCI-TEV-rv | TGCTTTCGCTGCCGCTTCTTTGGC |
| MacHis-TEV-fw | GCGGCGGCCAAAGCCGATTCTCACGAAGAACGCC |
| MacHis-TEV-rv | GGCGTTCTTCGTGAGAATCGGCTTTGGCCGCCGC |
| Plantaricin-TEV-fw | GCGAAAGAAGCGGCGGCCAAAGCCAAAAGCAGTGCGTATTCCTTGC |
| Plantaricin-TEV-rv | GGCTTTGGCCGCCGCTTCTTTCGC |
| Pleurocidin-TEV-fw | GCGAAAGAAGCGGCGGCCAAAGCCGGATGGGGCTCGTTCTTTAAGAAGGC |
| Pleurocidin-TEV-rv | GGCTTTGGCCGCCGCTTCTTTCGC |
| Magainin-TEV-fw | GCGAAAGAAGCGGCGGCCAAAGCCGGTATTGGGAAATTCCTGCATTCC |
| Magainin-TEV-rv | GGCTTTCGCTGCAGCCTCTTTCGC |
| MacHis-R6A-fw | TCTCACGAAGAAGCCCATCATGGTCGTC |
| MacHis-R6A-rv | GACGACCATGATGGGCTTCTTCGTGAG |
| MacHis-R10A-fw | CGCCATCATGGTGCCCATGGTCACCAC |
| MacHis-R10A-rv | GTGGTGACCATGGGCACCATGATGGCG |
| MacHis-Y16A-fw | GGTCACCACAAGGCCGGCCGCAAATTC |
| MacHis-Y16A-rv | GAATTTGCGGCCGGCCTTGTGGTGACC |
| MacHis-F20A-fw | GGCCGCAAAGCCCACGAGAAACATCACAGTCATCGTGGC |
| MacHis-F20A-rv | GCCACGATGACTGTGATGTTTCTCGTGGGCTTTGCGGCC |
| MacHis-R6A/R10A-fw | GAAGAAGCCCATCATGGTGCCCATGGTCACC |
| MacHis-R6A/R10A-rv | GGTGACCATGGGCACCATGATGGGCTTCTTC |
| MacHis-K15A-fw | GGTCACCACGCGTATGGCCGCAAATTCC |
| MacHis-K15A-rv | GGAATTTGCGGCCATACGCGTGGTGACC |

**Table S4.** Remaining fluorescence (in relative fluorescence units [RFU]) of the cuticular wax-coated MTP plate treated with buffer only ("buffer"), pure eGFP ("eGFP"), and the eGFP-MacHis constructs (wild-type $MH_{WT}$ and the variants $MH_{1\text{-}8}$) after initial treatment with buffer/eGFP/AP solution and removal of the supernatant (4 wells per system/variant for each of the 3 MTPs).

| MTP | well | buffer | eGFP | $MH_{WT}$ | $MH_1$ | $MH_2$ | $MH_3$ | $MH_4$ | $MH_5$ | $MH_6$ | $MH_7$ | $MH_8$ |
|---|---|---|---|---|---|---|---|---|---|---|---|---|
| 1 | 1 | 47 | 7638 | 11242 | 10738 | 10392 | 11404 | 10373 | 20489 | 12450 | 13595 | 11813 |
| | 2 | 45 | 7733 | 11423 | 11700 | 11118 | 11520 | 11808 | 20697 | 11630 | 13068 | 15629 |
| | 3 | 45 | 7563 | 11669 | 11406 | 10859 | 12756 | 12554 | 22576 | 11972 | 12097 | 11867 |
| | 4 | 52 | 8006 | 10183 | 10664 | 10831 | 11843 | 13303 | 25630 | 12752 | 14065 | 11351 |
| 2 | 1 | 47 | 10024 | 12834 | 12319 | 11387 | 11827 | 13513 | 23069 | 13557 | 12080 | 12380 |
| | 2 | 52 | 9322 | 14242 | 14231 | 11705 | 14132 | 14339 | 24713 | 13403 | 12843 | 12573 |
| | 3 | 48 | 9996 | 15348 | 14117 | 12400 | 15057 | 14269 | 24571 | 15493 | 13404 | 13072 |
| | 4 | 47 | 10547 | 14623 | 14638 | 12184 | 13802 | 15847 | 25559 | 15232 | 13884 | 13329 |
| 3 | 1 | 49 | 8556 | 11734 | 13707 | 12125 | 13055 | 14518 | 26681 | 13774 | 14949 | 14223 |
| | 2 | 50 | 9311 | 13282 | 13648 | 12538 | 13933 | 15496 | 26410 | 14627 | 14933 | 14625 |
| | 3 | 48 | 9092 | 13072 | 13222 | 11228 | 13560 | 14326 | 27762 | 15131 | 14908 | 16038 |
| | 4 | 47 | 9395 | 11716 | 12972 | 10934 | 13095 | 14302 | 25753 | 16248 | 16000 | 14197 |

**Table S5.** Remaining fluorescence (in relative fluorescence units [RFU]) of the cuticular wax-coated MTP plate treated with buffer only ("buffer"), pure eGFP ("eGFP"), and the eGFP-MacHis constructs (wild-type $MH_{WT}$ and the variants $MH_{1\text{-}8}$) after two washing steps (4 wells per system/variant for each of the 3 MTPs).

| MTP | well | buffer | eGFP | $MH_{WT}$ | $MH_1$ | $MH_2$ | $MH_3$ | $MH_4$ | $MH_5$ | $MH_6$ | $MH_7$ | $MH_8$ |
|---|---|---|---|---|---|---|---|---|---|---|---|---|
| 1 | 1 | 51 | 214 | 968 | 968 | 325 | 400 | 777 | 718 | 717 | 702 | 601 |
| | 2 | 51 | 212 | 976 | 962 | 305 | 435 | 782 | 752 | 714 | 713 | 598 |
| | 3 | 49 | 214 | 943 | 969 | 394 | 437 | 776 | 727 | 740 | 721 | 584 |
| | 4 | 51 | 225 | 1055 | 951 | 368 | 440 | 771 | 808 | 739 | 787 | 687 |
| 2 | 1 | 46 | 219 | 984 | 928 | 319 | 411 | 761 | 685 | 722 | 683 | 594 |
| | 2 | 50 | 210 | 969 | 877 | 400 | 389 | 745 | 753 | 685 | 674 | 591 |
| | 3 | 46 | 205 | 979 | 892 | 268 | 391 | 798 | 704 | 698 | 683 | 590 |
| | 4 | 47 | 211 | 997 | 901 | 314 | 398 | 761 | 764 | 725 | 734 | 615 |
| 3 | 1 | 45 | 202 | 998 | 899 | 265 | 350 | 746 | 666 | 671 | 703 | 542 |
| | 2 | 52 | 198 | 931 | 872 | 260 | 345 | 733 | 662 | 641 | 661 | 568 |
| | 3 | 43 | 197 | 939 | 868 | 264 | 352 | 742 | 692 | 648 | 709 | 589 |
| | 4 | 48 | 212 | 943 | 868 | 291 | 367 | 750 | 704 | 708 | 740 | 588 |

**Table S6.** Remaining fluorescence (in relative fluorescence units [RFU]) of the cuticular wax-coated MTP plate treated with buffer only ("buffer"), pure eGFP ("eGFP"), and the eGFP-MacHis constructs (wild-type $MH_{WT}$ and the variants $MH_{1-8}$) after five washing steps (4 wells per system/variant for each of the 3 MTPs).

| MTP | well | buffer | eGFP | $MH_{WT}$ | $MH_1$ | $MH_2$ | $MH_3$ | $MH_4$ | $MH_5$ | $MH_6$ | $MH_7$ | $MH_8$ |
|---|---|---|---|---|---|---|---|---|---|---|---|---|
| 1 | 1 | 50 | 190 | 886 | 877 | 270 | 350 | 715 | 665 | 675 | 632 | 530 |
| | 2 | 49 | 189 | 890 | 882 | 265 | 377 | 733 | 680 | 658 | 650 | 533 |
| | 3 | 46 | 195 | 869 | 885 | 312 | 352 | 716 | 663 | 690 | 662 | 520 |
| | 4 | 53 | 205 | 935 | 875 | 290 | 367 | 692 | 719 | 683 | 710 | 606 |
| 2 | 1 | 46 | 190 | 898 | 852 | 255 | 355 | 707 | 618 | 665 | 622 | 550 |
| | 2 | 51 | 195 | 878 | 802 | 338 | 339 | 693 | 697 | 627 | 632 | 528 |
| | 3 | 47 | 186 | 888 | 811 | 229 | 337 | 731 | 636 | 631 | 631 | 537 |
| | 4 | 49 | 186 | 901 | 823 | 274 | 344 | 678 | 689 | 658 | 680 | 536 |
| 3 | 1 | 48 | 192 | 906 | 836 | 230 | 318 | 704 | 619 | 633 | 634 | 497 |
| | 2 | 50 | 181 | 863 | 808 | 225 | 307 | 685 | 609 | 598 | 613 | 522 |
| | 3 | 49 | 185 | 861 | 799 | 239 | 314 | 684 | 626 | 582 | 656 | 540 |
| | 4 | 51 | 195 | 865 | 792 | 253 | 323 | 688 | 655 | 650 | 676 | 559 |

**Table S7.** Remaining fluorescence (in relative fluorescence units [RFU]) of the cuticular wax-coated MTP plate treated with buffer only ("buffer"), pure eGFP ("eGFP"), and the eGFP-APs (LCI, MacHis, Plantaricin, Pleurocidin, Magainin) after initial treatment with buffer/eGFP/eGFP-AP solution and removal of the supernatant (4 wells per system/variant for each of the 3 MTPs).

| MTP | well | buffer | eGFP | LCI | MacHis | Plantaricin | Pleurocidin | Magainin |
|---|---|---|---|---|---|---|---|---|
| 1 | 1 | 56 | 4766 | 4773 | 4445 | 6089 | 2965 | 28740 |
| | 2 | 56 | 4587 | 4798 | 4547 | 6498 | 2894 | 29813 |
| | 3 | 65 | 4768 | 4716 | 4503 | 6943 | 3067 | 31290 |
| | 4 | 57 | 5302 | 4835 | 4679 | 7382 | 3513 | 32198 |
| 2 | 1 | 59 | 4769 | 5787 | 4695 | 6853 | 3914 | 36062 |
| | 2 | 59 | 5240 | 6516 | 4602 | 7112 | 4083 | 39839 |
| | 3 | 72 | 5835 | 6115 | 5141 | 7109 | 4750 | 40479 |
| | 4 | 55 | 7709 | 6200 | 5067 | 8221 | 5016 | 41232 |
| 3 | 1 | 52 | 4058 | 4144 | 4733 | 5883 | 3006 | 25294 |
| | 2 | 53 | 4466 | 4617 | 4781 | 6506 | 3438 | 28197 |
| | 3 | 51 | 4728 | 5117 | 4954 | 7390 | 3572 | 29008 |
| | 4 | 53 | 5299 | 5916 | 4858 | 7051 | 3957 | 31875 |

**Table S8.** Remaining fluorescence (in relative fluorescence units [RFU]) of the cuticular wax-coated MTP plate treated with buffer only ("buffer"), pure eGFP ("eGFP"), and the eGFP-APs (LCI, MacHis, Plantaricin, Pleurocidin, Magainin) after two washing steps (4 wells per system/variant for each of the 3 MTPs).

| MTP | well | buffer | eGFP | LCI | MacHis | Plantaricin | Pleurocidin | Magainin |
|---|---|---|---|---|---|---|---|---|
| 1 | 1 | 56 | 573 | 1505 | 1120 | 1439 | 500 | 1171 |
| | 2 | 57 | 553 | 1345 | 1086 | 1488 | 431 | 1344 |
| | 3 | 59 | 568 | 1366 | 1174 | 1543 | 465 | 1298 |
| | 4 | 55 | 589 | 1354 | 1110 | 1680 | 494 | 1339 |
| 2 | 1 | 55 | 581 | 1385 | 1045 | 1379 | 465 | 1398 |
| | 2 | 61 | 560 | 1375 | 1182 | 1541 | 471 | 1455 |
| | 3 | 70 | 554 | 1382 | 1137 | 1391 | 555 | 1482 |
| | 4 | 55 | 640 | 1382 | 1242 | 1565 | 532 | 1436 |
| 3 | 1 | 54 | 549 | 1361 | 1074 | 1134 | 429 | 1515 |
| | 2 | 54 | 516 | 1383 | 1117 | 1460 | 429 | 1505 |
| | 3 | 49 | 509 | 1356 | 1092 | 1450 | 449 | 1439 |
| | 4 | 51 | 728 | 1356 | 1089 | 1160 | 501 | 1472 |

**Table S9.** Remaining fluorescence (in relative fluorescence units [RFU]) of the cuticular wax-coated MTP plate treated with buffer only ("buffer"), pure eGFP ("eGFP"), and the eGFP-APs (LCI, MacHis, Plantaricin, Pleurocidin, Magainin) after five washing steps (4 wells per system/variant for each of the 3 MTPs).

| MTP | well | buffer | eGFP | LCI | MacHis | Plantaricin | Pleurocidin | Magainin |
|---|---|---|---|---|---|---|---|---|
| 1 | 1 | 58 | 503 | 1494 | 1006 | 1166 | 424 | 1031 |
| | 2 | 59 | 484 | 1333 | 987 | 1197 | 372 | 1189 |
| | 3 | 60 | 498 | 1344 | 1063 | 1327 | 384 | 1182 |
| | 4 | 56 | 509 | 1354 | 1014 | 1396 | 437 | 1236 |
| 2 | 1 | 55 | 516 | 1358 | 966 | 1186 | 401 | 1262 |
| | 2 | 60 | 499 | 1369 | 1078 | 1309 | 407 | 1331 |
| | 3 | 64 | 482 | 1379 | 1035 | 1194 | 492 | 1353 |
| | 4 | 56 | 574 | 1378 | 1128 | 1435 | 445 | 1298 |
| 3 | 1 | 56 | 507 | 1370 | 968 | 940 | 383 | 1359 |
| | 2 | 53 | 465 | 1388 | 1028 | 1253 | 380 | 1399 |
| | 3 | 50 | 476 | 1363 | 1001 | 1175 | 398 | 1349 |
| | 4 | 55 | 636 | 1354 | 1006 | 1009 | 441 | 1377 |

### Supplementary Figures

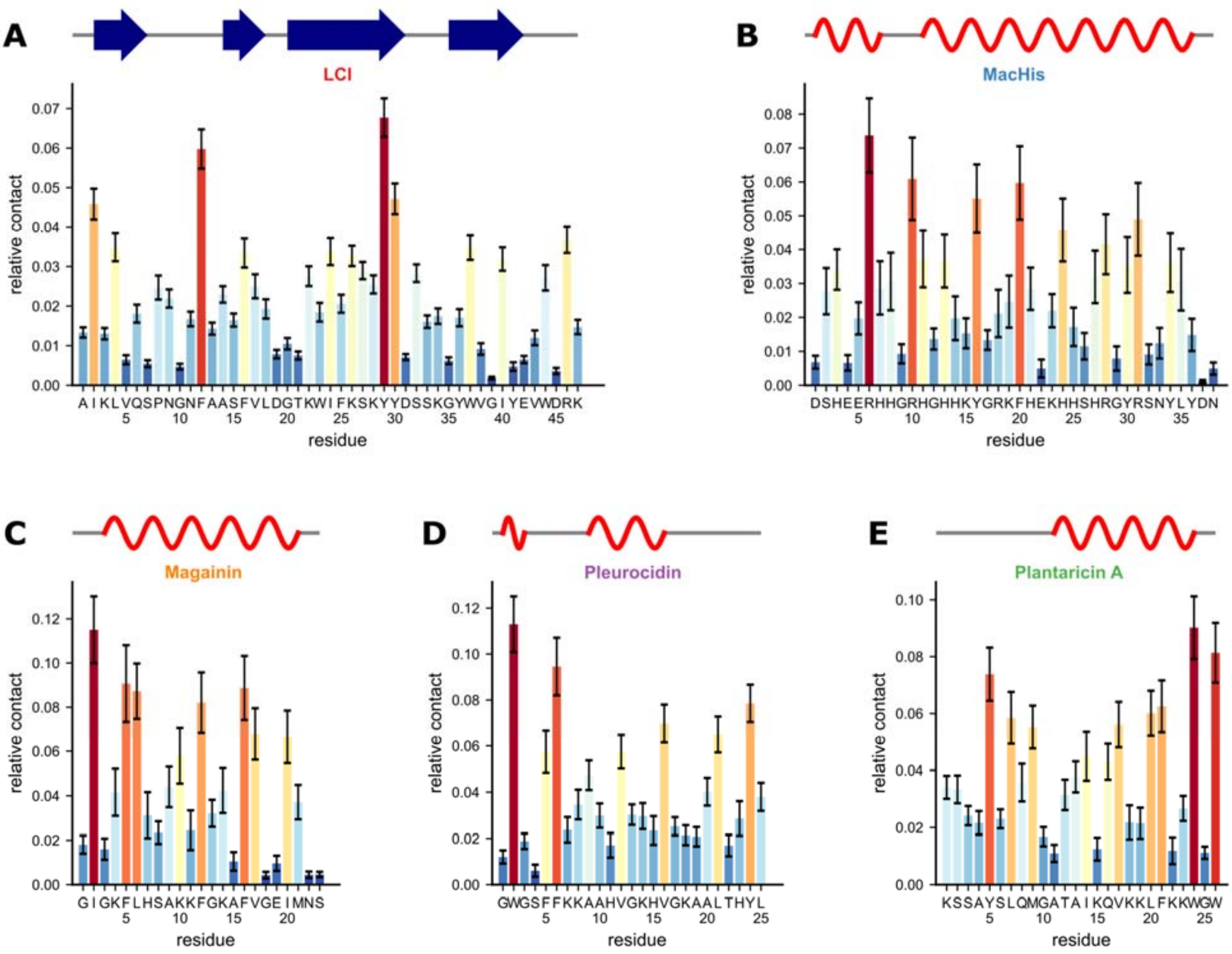

**Figure S1.** Residue-wise relative contacts of (A) LCI, (B) Macaque Histatin, (C) Magainin, (D) Pleurocidin, and (E) Plantaricin with the wax molecules within 10 x 250 ns long MD simulations of anchor peptide adsorption. Error bars denote the SEM. Geometrical analyses of the MD simulations were performed using cpptraj.[1] On top of the plots, the secondary structure of the APs is indicated (wavy line: α-helix, arrow: β-strand) as determined by DSSP[2] analysis of the protein structures.

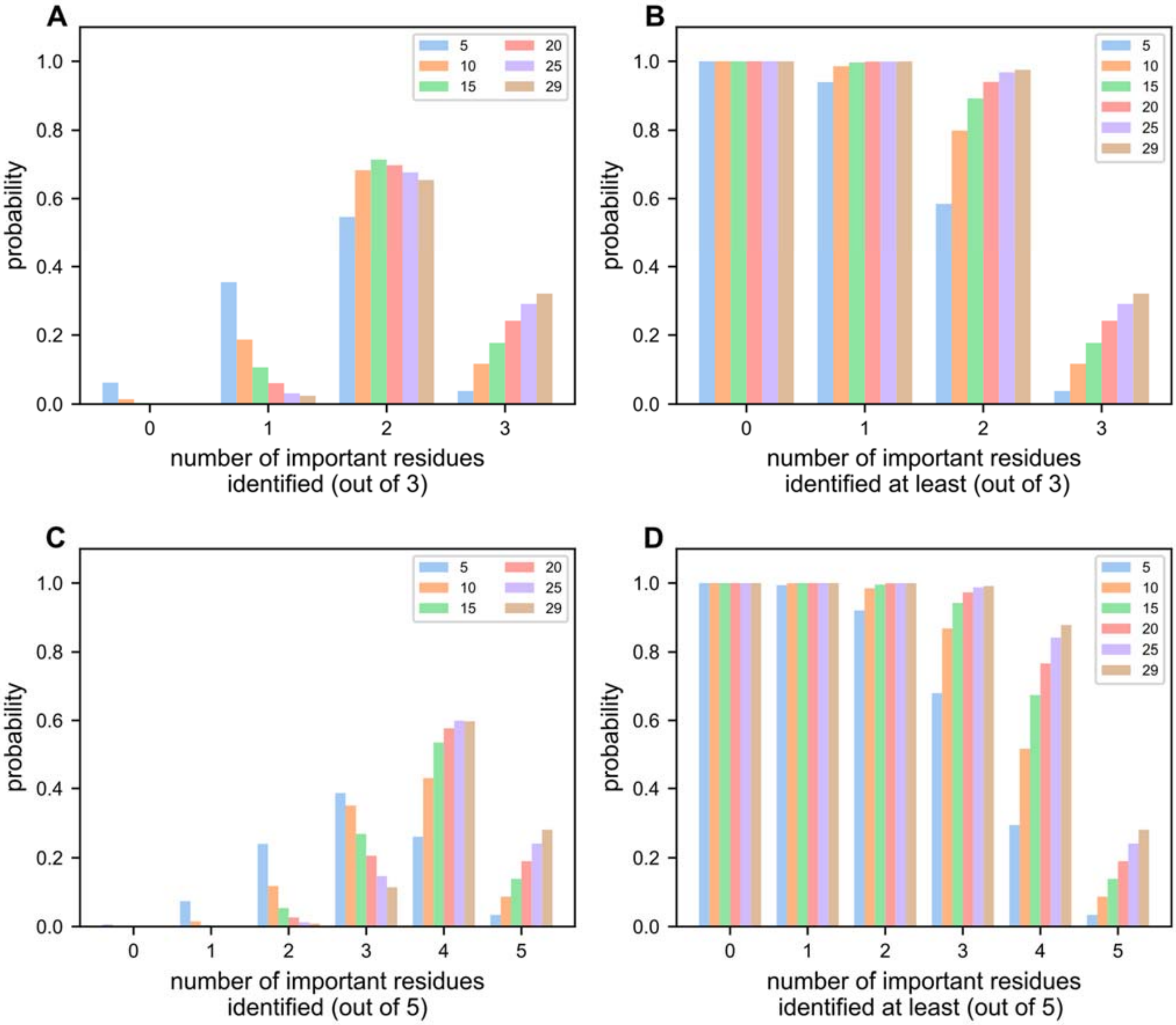

**Figure S2.** Bootstrapping analysis for identifying residues important for cuticular wax binding of MacHis in dependence of the sample size. Samples of different sizes (5, 10, 15, 20, 25, 29) were drawn 10,000 times with replacement from the original set containing 29 samples. For each sample set size, the overlap with the top three (**A**, **B**) and top five (**C**, **D**) identified residues using the original set is calculated. The probability of identifying exactly X residues out of the three/five is given in **A** and **C**, the probability of identifying at least X residues out of the three/five is given in **B** and **D** for each sample size.

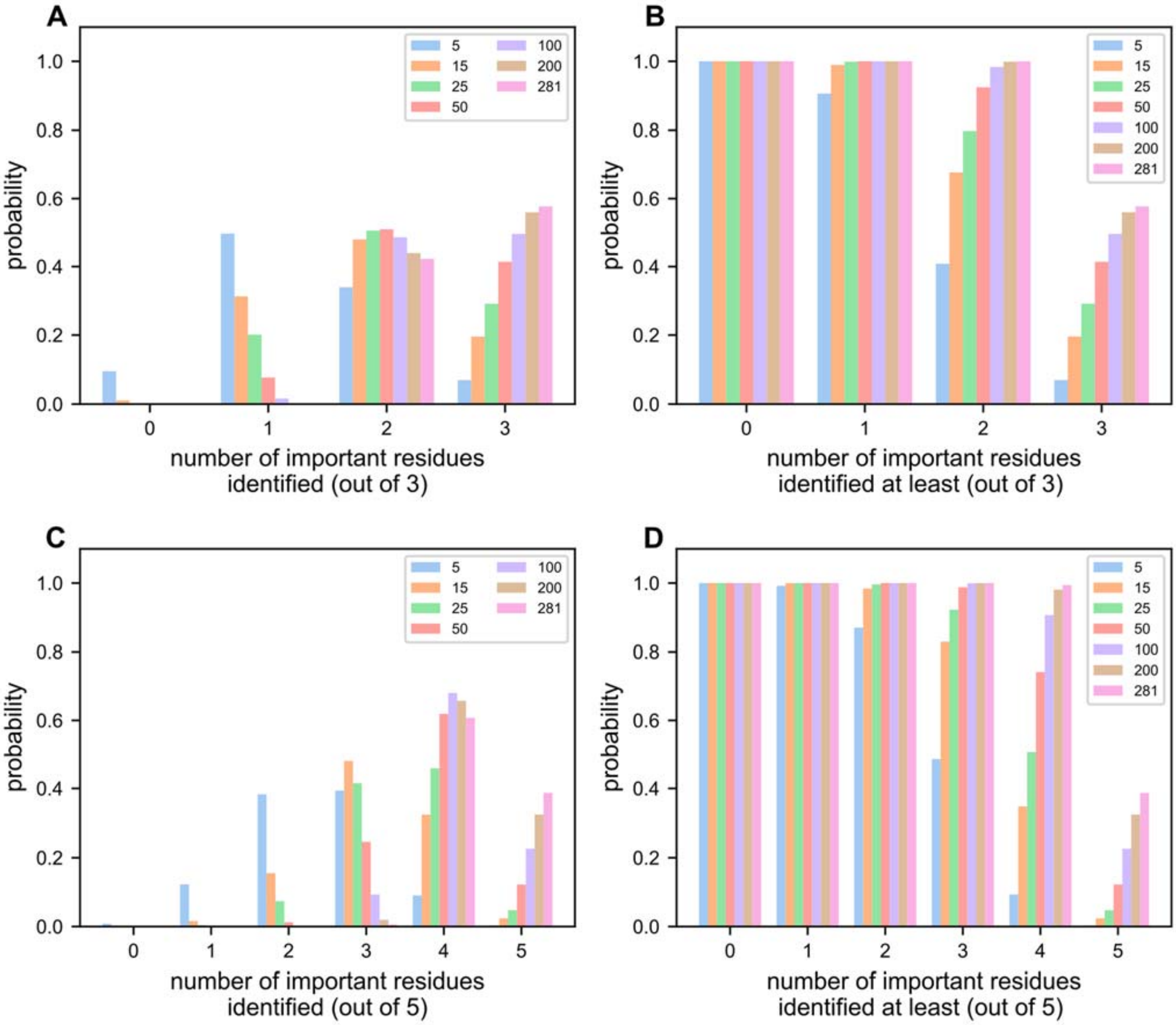

**Figure S3.** Bootstrapping analysis for identifying residues important for cuticular wax binding of LCI in dependence of the sample size. Samples of different sizes (5, 15, 25, 50, 100, 200, 281) were drawn 10,000 times with replacement from the original set containing 281 samples. For each sample set size, the overlap with the top three (**A**, **B**) and top five (**C**, **D**) identified residues using the original set is calculated. The probability of identifying exactly X residues out of the three/five is given in **A** and **C**, the probability of identifying at least X residues out of the three/five is given in **B** and **D** for each sample size.

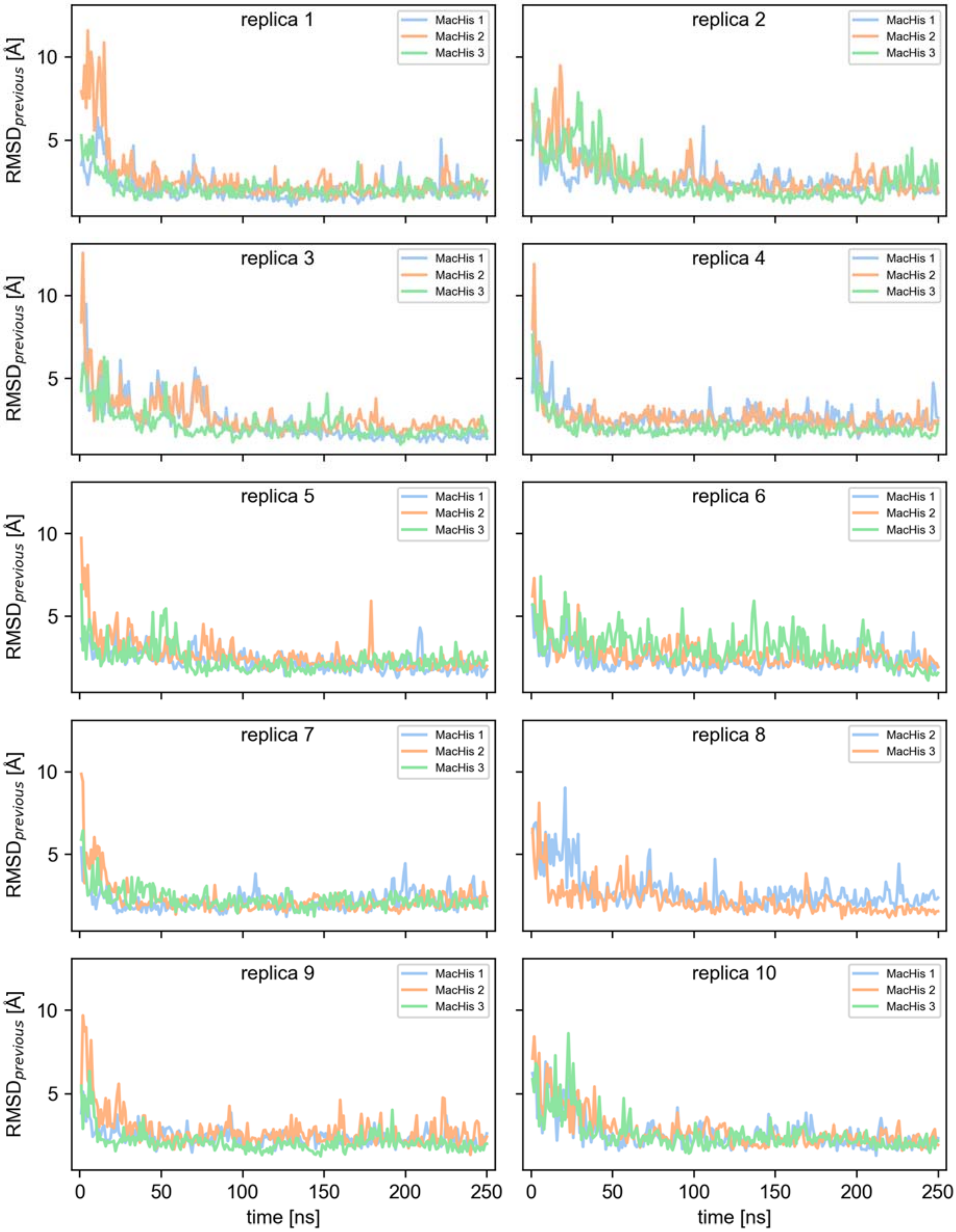

**Figure S4.** RMSD for all three MacHis peptides to their states 1 ns before, respectively, depicted for all ten simulated replicas. One peptide (MacHis 1, replica 8) is excluded, as it did not adsorb to the wax layer. Upon adsorption, observed within < 100 ns for the majority of the cases, the RMSD fluctuation of the peptide is decreased to 2.16 ± 0.04 Å (mean ± SEM for 100-250 ns)

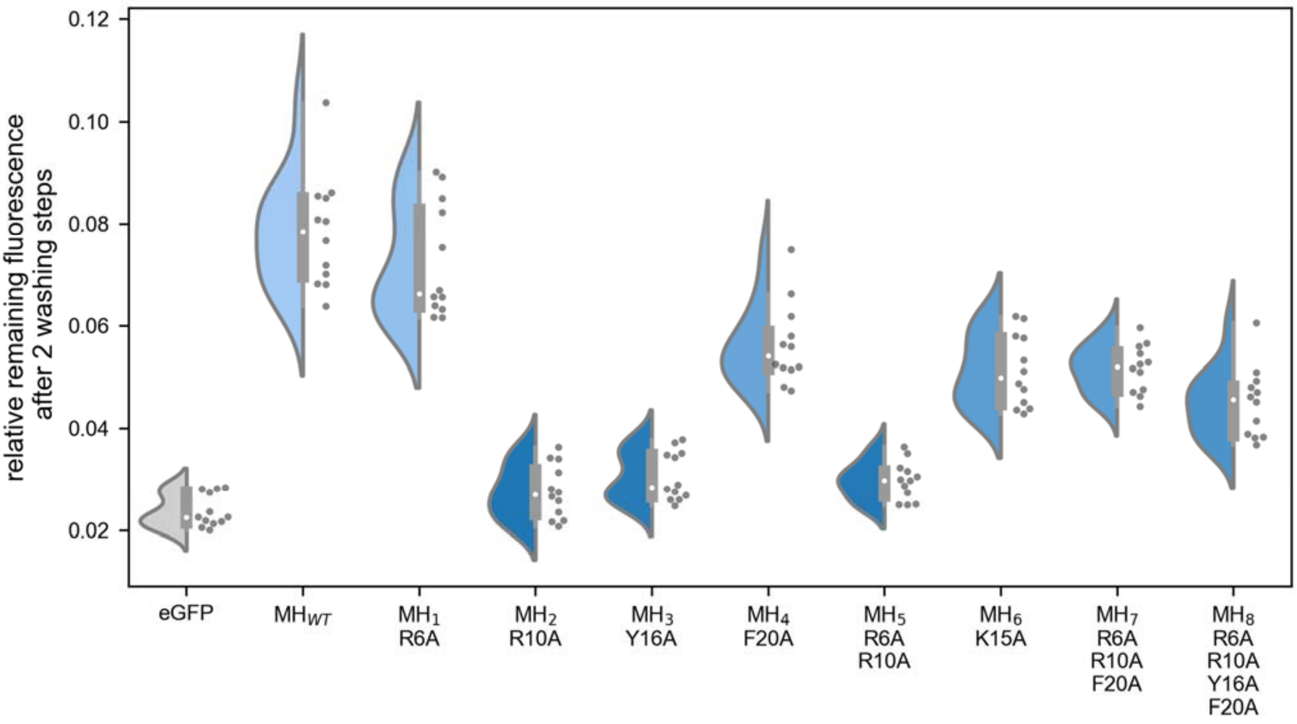

**Figure S5.** Binding of eGFP-MacHis wild-type (MH$_{WT}$) and eGFP-MacHis variants (MH$_{1\text{-}8}$) to extracted cuticular wax from apple leaves. Binding was quantified by measuring the remaining fluorescence of the fusion partner eGFP after five washing steps on apple leaf cuticular wax in 96-well-MTPs (4 wells, 3 replicates).

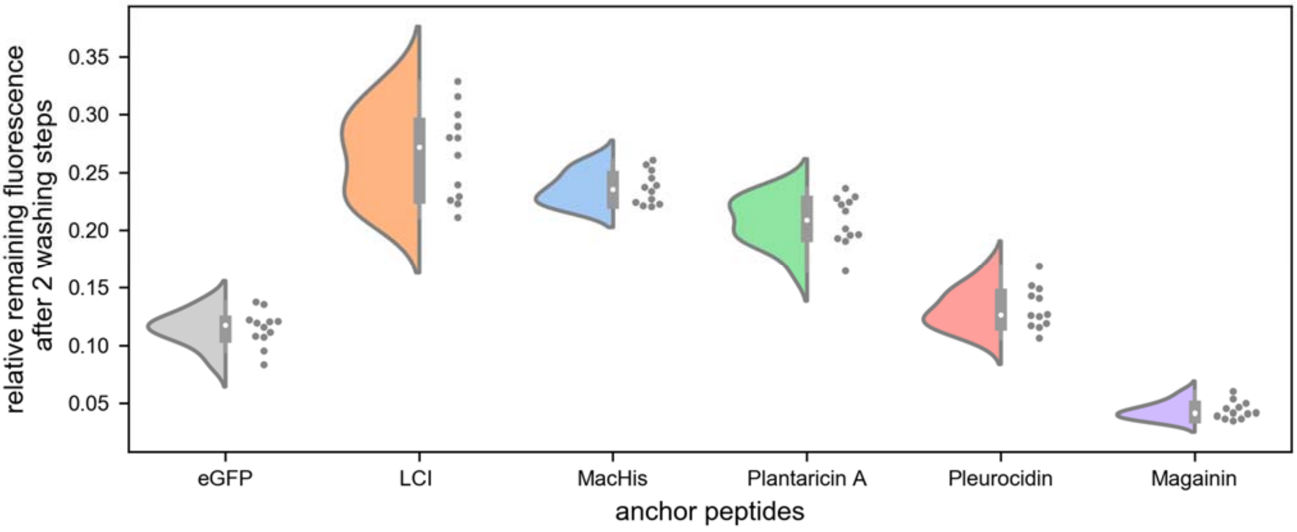

**Figure S6.** Binding of different eGFP-APs to extracted cuticular wax from apple leaves. Binding was quantified by measuring the remaining fluorescence of the fusion partner eGFP after two washing steps on apple leaf cuticular wax in 96-well-MTPs (4 wells, 3 replicates).

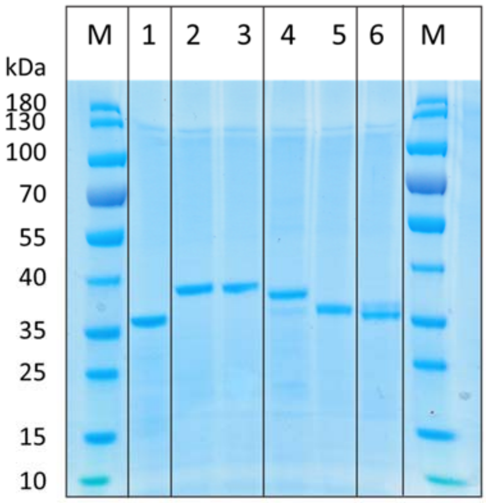

| | | |
|---|---|---|
| 1 | eGFP 1 µg | 31.5 kDa |
| 2 | LCI no TEV 1 µg | 36.5 kDa |
| 3 | MacHis no TEV 1 µg | 35.4 kDa |
| 4 | Plantaricin A no TEV 1 µg | 33.6 kDa |
| 5 | Pleurocidin no TEV 1 µg | 33.4 kDa |
| 6 | Magainin 2 no TEV 1 µg | 33.1 kDa |

**Figure S7.** Sodium dodecyl sulfate-polyacrylamide gel electrophoresis (SDS-PAGE) image of eGFP and eGFP-APs for determination of purity with ImageJ[3] (version 1.53e).

### Supplementary References

(1) Roe, D. R.; Cheatham, T. E., PTRAJ and CPPTRAJ: Software for Processing and Analysis of Molecular Dynamics Trajectory Data. *J. Chem. Theory Comput.* **2013,** *9* (7), 3084-3095.

(2) Kabsch, W.; Sander, C., Dictionary of protein secondary structure: Pattern recognition of hydrogen-bonded and geometrical features. *Biopolymers* **1983,** *22* (12), 2577-2637.

(3) Rasband, W., ImageJ: Image processing and analysis in Java. *Astrophysics Source Code Library* **2012**, ascl: 1206.013.
